# Adjudicating between face-coding models with individual-face fMRI responses

**DOI:** 10.1101/029603

**Authors:** Johan D. Carlin, Nikolaus Kriegeskorte

## Abstract

The perceptual representation of individual faces is often explained with reference to a norm-based face space. In such spaces, individuals are encoded as vectors where identity is primarily conveyed by direction and distinctiveness by eccentricity. Here we measured human fMRI responses and psychophysical similarity judgments of individual face exemplars, which were generated as realistic 3D animations using a computer-graphics model. We developed and evaluated multiple neurobiologically plausible computational models, each of which predicts a representational distance matrix and a regional-mean activation profile for 24 face stimuli. In the fusiform face area, a face-space coding model with sigmoidal ramp tuning provided a better account of the data than one based on exemplar tuning. However, an image-processing model with weighted banks of Gabor filters performed similarly. Accounting for the data required the inclusion of a measurement-level population averaging mechanism that approximates how fMRI voxels locally average distinct neuronal tunings. Our study demonstrates the importance of comparing multiple models and of modeling the measurement process in computational neuroimaging.

**Author Summary:** Humans recognize conspecifics by their faces. Understanding how faces are recognized is an open computational problem with relevance to theories of perception, social cognition, and the engineering of computer vision systems. Here we measured brain activity with functional MRI while human participants viewed individual faces. We developed multiple computational models inspired by known response preferences of single neurons in the primate visual cortex. We then compared these neuronal models to patterns of brain activity corresponding to individual faces. The data were consistent with a model where neurons respond to directions in a high-dimensional space of faces. It also proved essential to model how functional MRI voxels locally average the responses of tens of thousands of neurons. The study highlights the challenges in adjudicating between alternative computational theories of visual information processing.

## Introduction

Humans are expert at recognizing individual faces, but the mechanisms that support this ability are poorly understood. Multiple areas in human occipital and temporal cortex exhibit representations that distinguish individual faces, as indicated by successful decoding of face identity from functional magnetic resonance imaging (fMRI) response patterns (1–10). Decoding can reveal the presence of face-identity information as well as invariances. However, the nature of these representations remains obscure because individual faces differ along many stimulus dimensions, each of which could plausibly support decoding. To understand the representational space, we need to formulate models of how individual faces might be encoded and test these models with responses to sufficiently large sets of face exemplars. Here we use representational similarity analysis (RSA) (11) to test face-coding models at the level of the representational distance matrices they predict. Comparing models to data in the common currency of the distance matrix enables us to pool the evidence over many voxels within a region, obviating the need to fit models separately to noisy individual fMRI voxels.

Many cognitive and neuroscientific models of face processing do not make quantitative predictions about the representation of particular faces (12,13). However, such predictions can be obtained from models based on the notion that faces are encoded as vectors in a space (14). Most face-space implementations apply principal components analysis (PCA) to face images or laser scans in order to obtain a space, where each component is a dimension and the average face for the training sample is located at the origin (15,16). In such PCA face spaces, eccentricity is associated with judgments of distinctiveness, while vector direction is associated with perceived identity (17–19). Initial evidence from macaque single-unit recordings and human fMRI suggests that brain responses to faces are strongly modulated by face-space eccentricity, with most studies finding increasing responses with distinctiveness (20–23). However, there has been no attempt to develop a unified account for how a single underlying face-space representation can support both sensitivity to face-space direction at the level of multivariate response patterns and sensitivity to eccentricity at the level of regional-mean fMRI activations. Here we develop and evaluate several face-space coding models, which differ with respect to the proposed shape of the neuronal tuning functions across face space and with respect to the distribution of preferred face-space locations over the simulated neuronal population.

Face-space coding models define high-level representational spaces, which are assumed to arise through unspecified low-level featural processing. An alternative possibility is that some cortical face-space representations can be explained directly by low-level visual features, which typically covary with position in PCA-derived face spaces. To explore this possibility, we evaluated a Gabor-filter model, which receives stimulus images rather than face-space coordinates as input, and has previously been used to model response preferences of individual voxels in early visual cortex (24).

We found that cortical face responses measured with fMRI strongly reflect face-space eccentricity: a step along the radial axis in face space results in a much larger pattern change than an equal step along the tangential axis. These effects were consistent with either a sigmoidal-ramp-tuning or a Gabor-filter model. The performance of these winning models depended on the inclusion of a measurement-level population-averaging mechanism, which accounted for local averaging of neuronal tunings in fMRI voxel measurements.

## Results

### Sampling face space with photorealistic but physically-controlled animations

In order to elicit strong percepts of the 3D shape of each individual face, we generated a set of photorealistic animations of face exemplars. Each 2s animation in the main experiment featured a face exemplar in left or right half profile, which rotated outward continuously (S2 Movie, Materials and Methods). The animations were based on a PCA model of 3D face shape and texture (25). Each frame of each animation was cropped with a feathered aperture and processed to equate low-level image properties across the stimulus set (Materials and Methods). We generated 12 faces from a slice through the PCA face space (Fig 1b). Euclidean distances between the Cartesian coordinates for each face were summarized in a distance matrix (Fig 1a), which served as the reference for comparisons against distances in the perceptual and cortical face spaces (Fig 1c-h). We generated a physically distinct stimulus set with the same underlying similarity structure for each participant by randomizing the orientation of the slice through the high-dimensional PCA space (for examples, see S1 Fig). This served to improve generalizability by ensuring that group-level effects were not strongly influenced by idiosyncrasies of face exemplars drawn from a particular PCA space slice. In formal terms, group-level inference therefore treats stimulus as a random effect (26).

**Fig 1.**
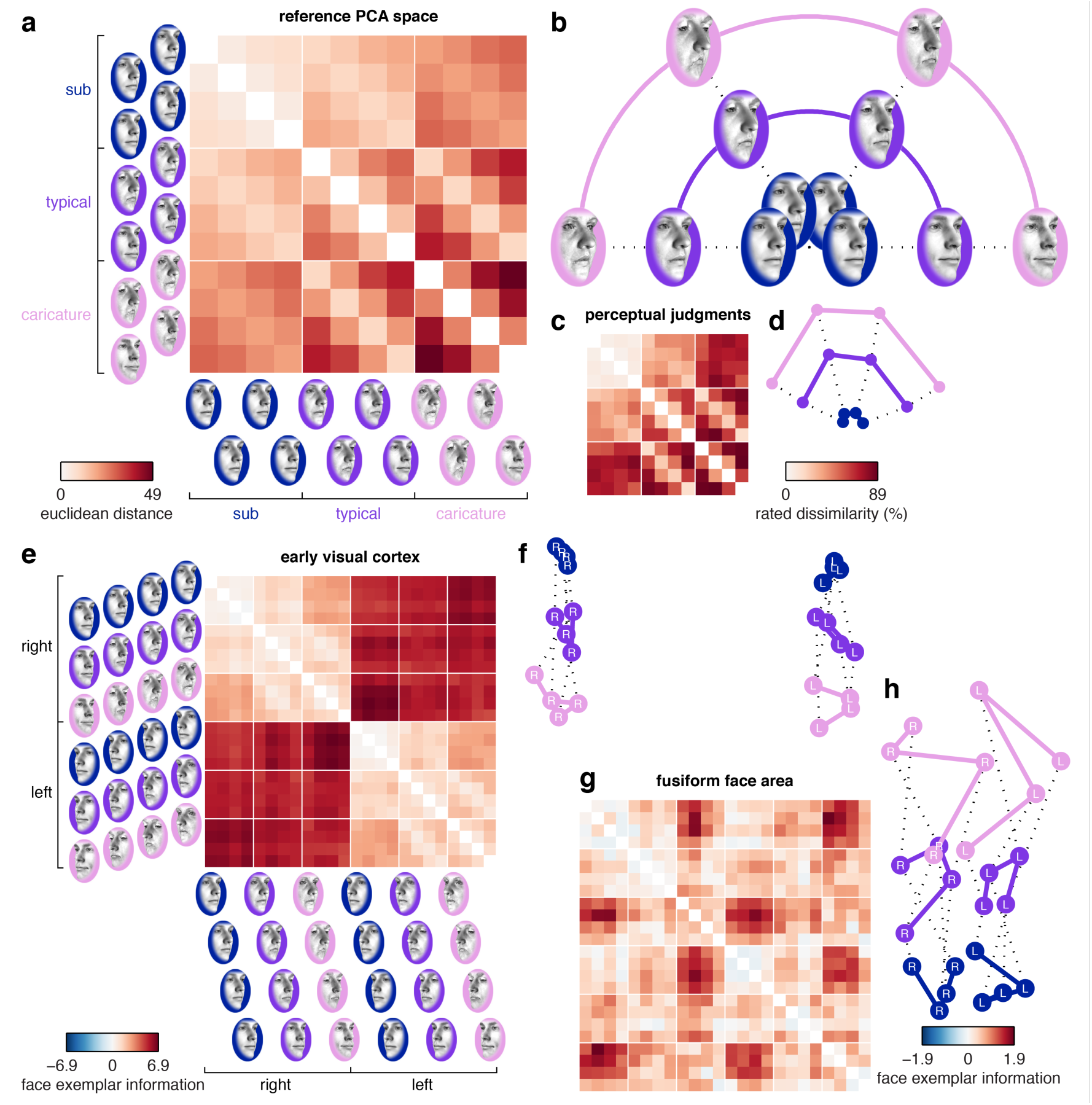
Face spaces obtained from a reference PCA model, perceptual similarity judgments and fMRI response patterns in human visual cortex. (**a-b**) We generated 12 faces in a polar grid arrangement on a 2D slice through the reference PCA space (3 eccentricity levels and 4 directions). The grid was centered on the stimulus space norm (not shown). Euclidean distances between the 12 faces are illustrated in a distance matrix (a) and in a 2D visualization of these distances (b, multidimensional scaling, metric-stress criterion). (**c-d**) Group-average perceptual face space (N=10) estimated from psychophysical similarity judgments. The distance matrix depicts the percentage of trials on which each pair of faces was rated as relatively more dissimilar. (**e-h**) Group-average cortical face-space (N=10) estimated from fMRI response patterns in human visual cortex. The distance matrices depict a cross-validated estimate of multivariate discriminability where 0 corresponds to chance-level performance (leave-one-run-out cross-validation, Materials and Methods). Each cortical region was defined in individual participants using independent data. The distance matrices use separate color scales since we expect variable effect sizes across cortical regions. For data from other regions of interest, see S2 Fig.

### Cortical face spaces are warped relative to the reference PCA space

In order to sample human cortical and perceptual face representations, human participants (N=10) participated in a perceptual judgment task followed by fMRI scans. The perceptual judgment task (S1 Movie, 2145 trials over 4 recording days) involved force-choice judgments of the relative similarity between pairs of faces, which were used to estimate a behavioral distance matrix (Materials and Methods). The same faces were then presented individually in a subsequent rapid event-related fMRI experiment (S2 Movie, 2496 trials over 4 recording days). Brain responses were analyzed separately in multiple independently localized visual regions of interest. We compared representational distance matrices estimated from these data sources to distances predicted according to different models using the Pearson correlation coefficient. These distance-matrix similarities were estimated in single participants and the resulting coefficients were Z-transformed and entered into a summary-statistic group analysis for random-effects inference generalizing across participants and stimuli (Materials and Methods).

We observed a strong group-average correlation between distance matrices estimated from perceptual dissimilarity judgments and Euclidean distances in the reference PCA space (mean(r) = 0.83, mean(Z(r)) = 1.20, standard error = 0.05, p < 0.001, Fig 1c, Fig 5a, S1 Table). Correlations between the reference PCA space and cortical face spaces were generally statistically significant, but smaller in magnitude (Fig 5b-c, S5 Fig, S1 Table). Distances estimated from the fusiform face area were weakly, but highly significantly correlated with the reference PCA space (mean(r) = 0.17, mean(Z(r)) = 0.17, standard error = 0.04, p < 0.001, Fig 5c). Distances estimated from the early visual cortex were even less, though still significantly, correlated with the reference PCA space (mean(r) = 0.07, mean(Z(r)) = 0.07, standard error = 0.04, p = 0.044, Fig 5b). These smaller correlations in cortical compared to perceptual face spaces could not be attributed solely to lower functional contrast-to-noise ratios in fMRI data, because the effects generally did not approach the noise-ceiling estimate for the sample (shaded region in Fig 5). The noise ceiling was based on the reproducibility of distance matrices between participants (Materials and Methods). Instead, these findings indicate that the reference PCA space could not capture all the explainable dissimilarity variance in cortical face spaces.

### Cortical face spaces over-represent eccentricity

We quantified the apparent warps in the cortical face spaces by constructing a multiple-regression RSA model, with separate distance-matrix predictors for eccentricity and direction, and for within and across face viewpoints (Fig 2a, Materials and Methods). These predictors were scaled such that differences between the eccentricity and direction parameters could be interpreted as warping relative to a veridical encoding of distances in the reference PCA face space. We also observed strong viewpoint effects in multiple regions, including the early visual cortex (Fig 1e-f). Such effects were modeled by separate constant terms for distances within and across viewpoint.

**Fig 2.**
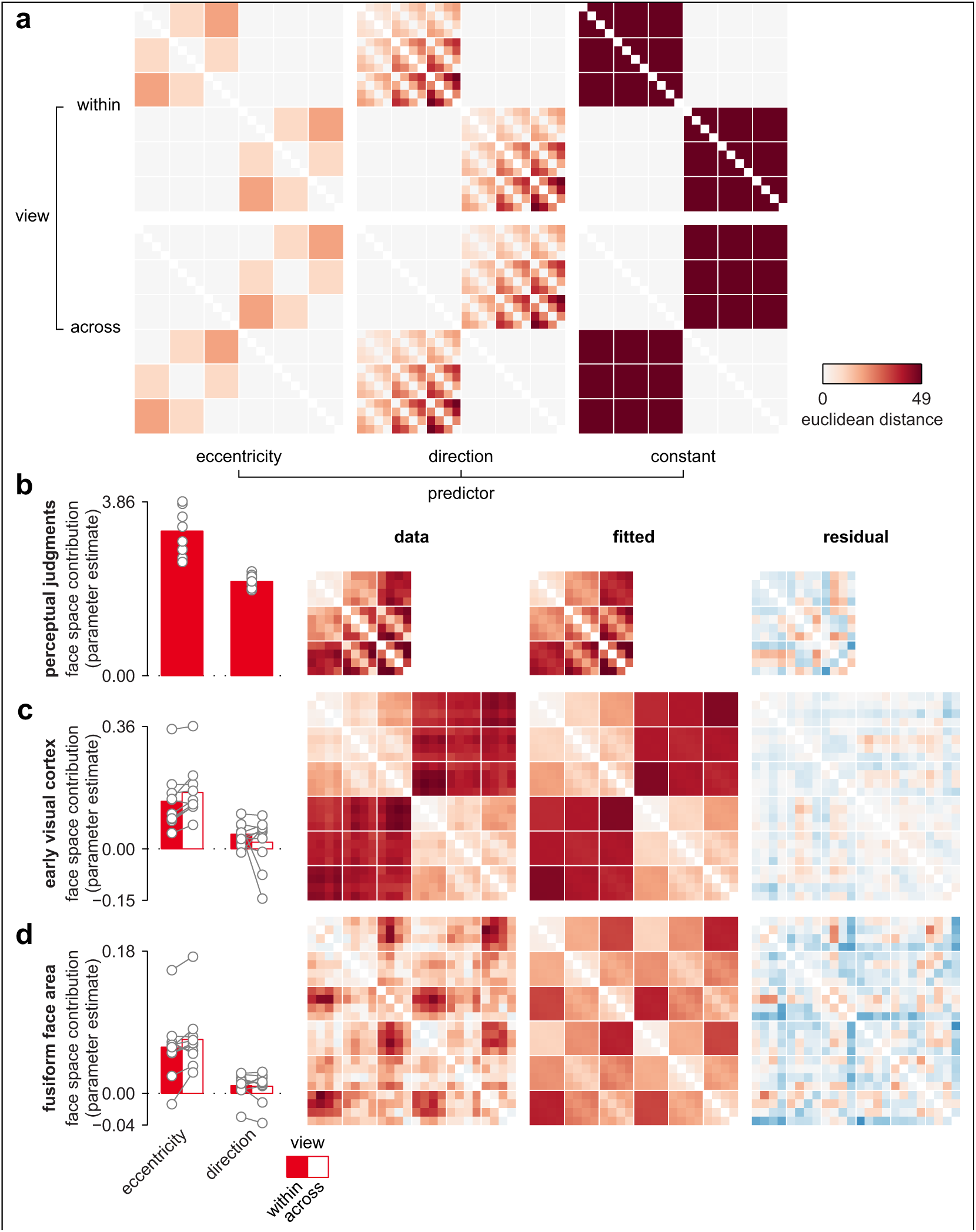
Face-space warping reflects an over-representation of eccentricity over direction information. (**a**) Squared distances in the reference PCA space were parameterized into predictors coding all 4 combinations of face-space metric (direction, eccentricity) and viewpoint (within, across). This multiple regression RSA model also included separate constant terms for each viewpoint, and was fitted separately to each face space using ordinary least squares (Materials and Methods). Importantly, the scaling of the eccentricity and direction predictors ensures that equal parameter estimates corresponds to a preserved reference PCA space. (**b**) Multiple regression fit to the perceptual face space with group-average parameter estimates, distance matrix, fitted distances and residuals. Gray lines reflect single participant parameter estimates (**c-d**) Multiple regression fit to the cortical face spaces, plotted as in b. See S2 Table for inferential statistics.

Eccentricity changes had a consistently greater effect on representational patterns than direction changes, suggesting that a step change along the radial axis resulted in a larger pattern change than an equivalent step along the tangential axis (Fig 2c-d). The overrepresentation of eccentricity relative to direction was observed in each participant, and in both cortical and perceptual face spaces, although the effect was considerably larger in cortical face spaces. A repeated-measures two-factor ANOVA (eccentricity versus direction, within versus across viewpoint) on the single-participant parameter estimates from the multiple-regression RSA model was consistent with these apparent differences, with a statistically significant main effect of eccentricity versus direction for perceptual and cortical face spaces (all p<0.012, S2 Table). Thus, compared to the encoding in the reference PCA space, cortical face spaces over-represented the radial, distinctiveness-related axis compared to the tangential, identity-related axis.

Although these findings suggest a larger contribution of face-space eccentricity than direction in visual cortex, we also observed clear evidence for greater-than-chance discrimination performance among faces that differed only in face-space direction. Group-average cross-validated discriminant distances for faces that differed in direction but not eccentricity exceeded chance-level performance (p < 0.05) for typical and caricatured faces in both the early visual cortex and the fusiform face area (S3 Fig). Sub-caricatured faces were less consistently discriminable. Indeed, direction discrimination increased with eccentricity both within (mean = 0.013, standard error = 0.007, p = 0.036) and across (mean = 0.012, standard error = 0.005, p = 0.016) viewpoint in the fusiform face area (linear effect of sub > typical > caricature), suggesting that direction discrimination increased with face-space eccentricity in a dose-dependent manner (S3 Table). By contrast, within viewpoint discrimination performance in the early visual cortex scaled with eccentricity (mean = 0.039, standard error = 0.008, p < 0.001), but distances that spanned a viewpoint change did not vary with eccentricity (mean = 0.009, standard error = 0.019, p = 0.297). This is consistent with a view-dependent representation in early visual cortex. Thus, cortical regions discriminate identity-related direction information even in the absence of a difference in distinctiveness-related eccentricity information, suggesting that cortical face representations cannot be reduced to a one-dimensional code based on distinctiveness alone. In summary, cortical coding of face-space position is systematically warped relative to the reference PCA space, with a substantial overrepresentation of eccentricity and a smaller, but reliable contribution of face-space direction.

### Regional-mean fMRI activation increases with face-space eccentricity, but removing such effects does not substantially alter cortical face-space warping

Cortical face-space warping could not be explained by regional-mean activation preferences for caricatures. We performed a regional-mean analysis of responses in each cortical area, which confirmed previous reports that fMRI responses increase with distinctiveness across much of visual cortex (Fig 6, S6 Fig) (23). In order to test the influence of such regional-mean activation effects on representational distances, we adapted our discriminant distance metric to remove additive and multiplicative overall activation effects (Materials and Methods). Distance matrices estimated using this alternative method were highly similar to ones estimated without removal of overall activation effects (all r = 0.9 or greater for the Pearson correlation between group-average distance matrices with and without mean removal, S4 Fig). Thus, although eccentricity affected the overall activation in all visual areas, the warping of the cortical face spaces could not be attributed to overall activation effects alone.

### Accounting for cortical and perceptual face spaces with PCA face-space-tuning models and image-computable models

We developed multiple computational models, each of which predicts a representational distance matrix and a regional-mean activation profile. These models can be divided into three classes: the sigmoidal-ramp tuning and exemplar tuning models receive face-space coordinates as input, while the Gabor-filter model receives gray-scale pixel intensities from the stimulus images as input. We evaluated each of these three model classes with and without a measurement-level population-averaging mechanism, which approximates how fMRI voxels locally average underlying neural activity.

The sigmoidal-ramp-tuning model proposes that the representational space is covered with randomly oriented ramps, each of which exhibits a monotonically increasing response along its preferred direction in face space (Fig 3a). This model is inspired by known preferences for extreme feature values in single units recorded from area V4 and from face-selective patches in the macaque visual cortex (20,27,28). We modeled the response along each model neuron’s preferred direction using a sigmoidal function with two free parameters, which control the horizontal offset and the saturation of the response function (Materials and Methods). A third parameter controlled the strength of measurement-level population averaging by translating each individual model unit’s response toward the population-mean response. The way this accounts for local averaging by voxels is illustrated for the fusiform face area in Fig 3a-b. It can be seen that measurement-level population averaging introduces a substantial U-shape in the individual response functions, with only a minor deflection in favor of a preferred face-space direction. At the level of Euclidean distances between population response vectors evoked by each face, this leads to exaggerated distances for radial relative to tangential face differences (Fig 3c). In summary, measurement-level population averaging provides a simple means to interpolate between two extreme cases: A value of 0 corresponds to the case where the model’s response is perfectly preserved in the fMRI voxels, whereas a value of 1 corresponds to the case where the model’s response to a given stimulus is reduced to the arithmetic mean over the model units.

**Fig 3.**
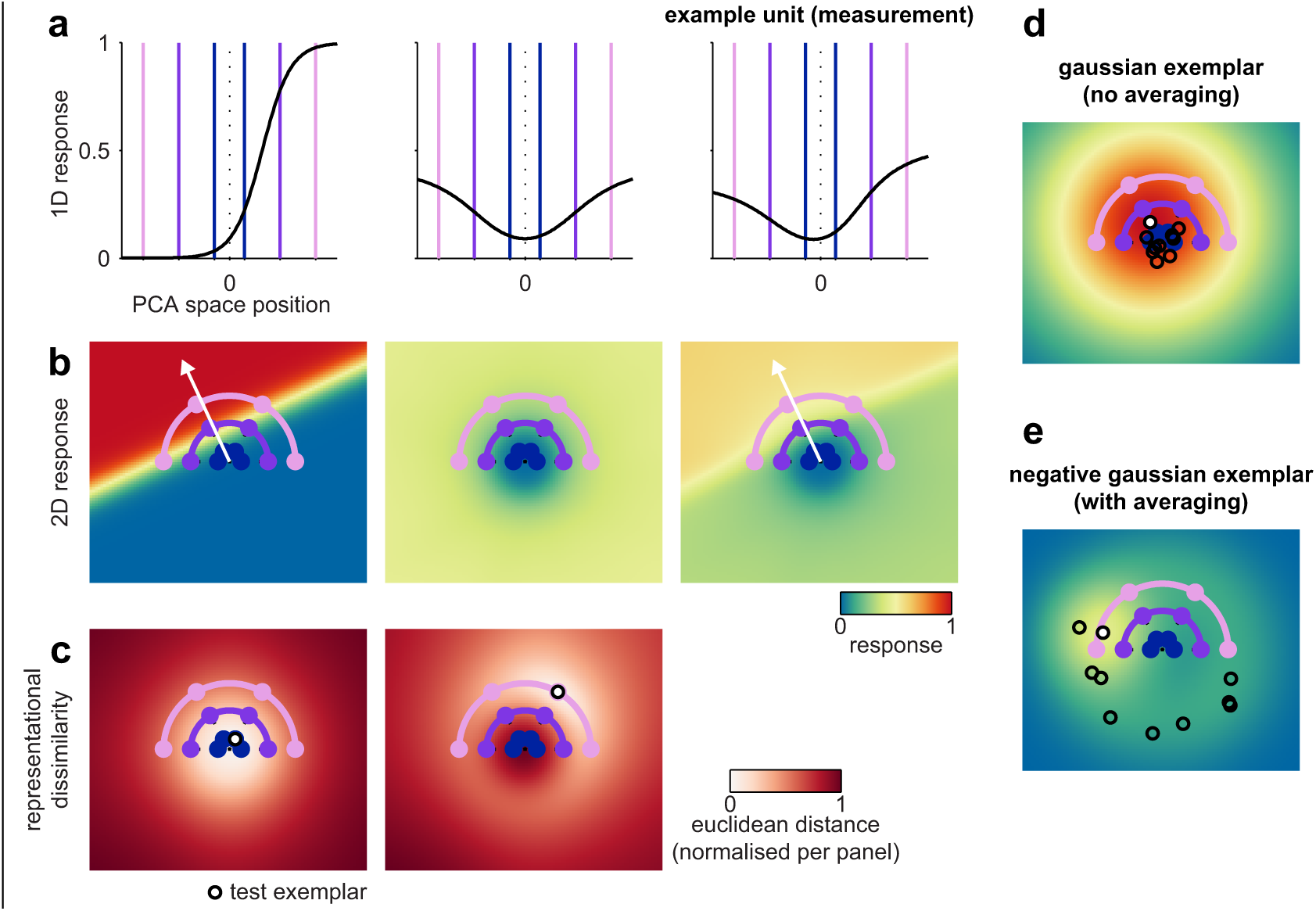
Computational models for face-space coding based on sigmoidal ramp or exemplar tuning coupled with measurement-level population averaging. The visualization uses the model parameters that were optimal for predicting the face space in the fusiform face area. (**a**) Response function from the sigmoidal ramp model’s internal representation (left panel) and its measurement (right panel) following translation toward the population-average response function (middle panel). (**b**) Two-dimensional generalization of the sigmoidal ramp tuning function to encode a direction in the PCA-space slice. An example unit is plotted in the left and right panels with the population-average response in the middle panel. (**c**) The model’s representational dissimilarity structure was estimated as the Euclidean distance between the population response vectors elicited by a coordinate in the PCA-space slice (white circle) and every other coordinate on the face space slice. Two example coordinates are plotted in the left and right panels. It can be seen that dissimilarity increases more rapidly with radial (eccentricity) than with tangential (direction) face-space distance. (**d**) Two-dimensional response function for an example unit from the Gaussian exemplar model (white marker). A sub-set of other units is overlaid in black markers to illustrate the width of the Gaussian distribution of tuning centers. (**e**) Two-dimensional response function for an example unit from the negative Gaussian exemplar model, plotted as in panel d.

In the exemplar model, each unit prefers a location in face space, rather than a direction, and its tuning is described by a Gaussian centered on the preferred location. The representational space is covered by a population of units whose preferred locations are sampled from a Gaussian centered on the norm face (Fig 3d, Materials and Methods). We fitted the Gaussian exemplar-tuning model similarly to the sigmoidal ramp-tuning model, using two parameters that controlled the width of the Gaussian tuning function and the width of the Gaussian distribution from which preferred faces were sampled. We also evaluated a variant of the exemplar model where the distribution of preferred faces followed an inverted-Gaussian distribution (Fig 3e). Population averaging was modeled in the same way as for the sigmoidal-ramp-tuning model using a third parameter.

The Gabor-filter model differs from the previous model classes in that it receives gray-scale image intensities as input, rather than PCA space position (Fig 4a). Such models have previously been used to account for response preferences of individual voxels in early visual cortex (24). The model comprises Gabor filters varying in orientation, spatial frequency and phase, and spatial position. The filters are organized into banks, each corresponding to a spatial frequency and comprising a different number of spatial positions (coarser for lower spatial frequencies; Fig 4b, Materials and Methods). We assumed that all orientations and spatial positions are equally represented. For the spatial frequencies, however, we let the data determine the weighting. We fitted a weighted representational model with one weight for each spatial-frequency bank (5 free parameters, Fig 4c). Local averaging in fMRI voxels was modeled using two stages of measurement-level population averaging: First, filters with tuning centers on either side of the vertical meridian were translated separately toward their respective hemifield-specific population averages. Second, a global-pool averaging was performed similarly to the other models. The contribution of these two population-average signals to the measured responses was modeled by 2 additional parameters (Fig 4d). The additional hemifield-specific averaging stage was necessary to account for strong view-specific effects in early visual cortex, but did not materially contribute to the fit in ventral temporal regions.

**Fig 4.**
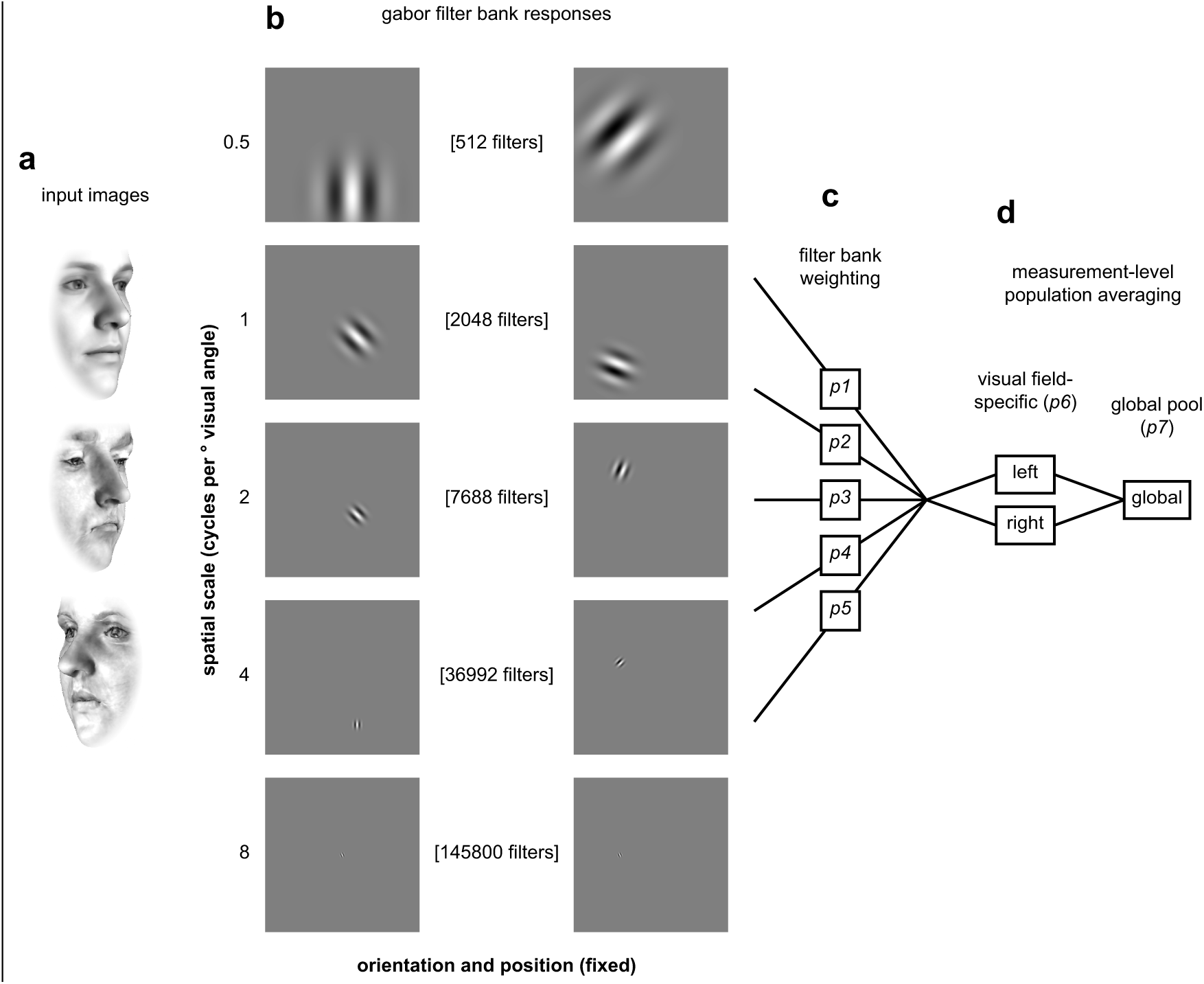
Schematic illustration of processing stages for the Gabor filter model. (**a**) The model receives gray-scale image intensities for each face exemplar. Three examples input images are illustrated in rows. (**b**) Image intensities are passed through banks of Gabor filters. The banks vary in spatial scale (filter standard deviation and grid spacing). The rows illustrate example filters from each bank. (**c**) The output of each filter bank is weighted and measurement effects are modeled using subsequent hemifield-specific and global-pool population averaging stages (**d**). The final output of the model is a Euclidean distance matrix estimated from the activation vectors in response to each face.

**Fig 5.**
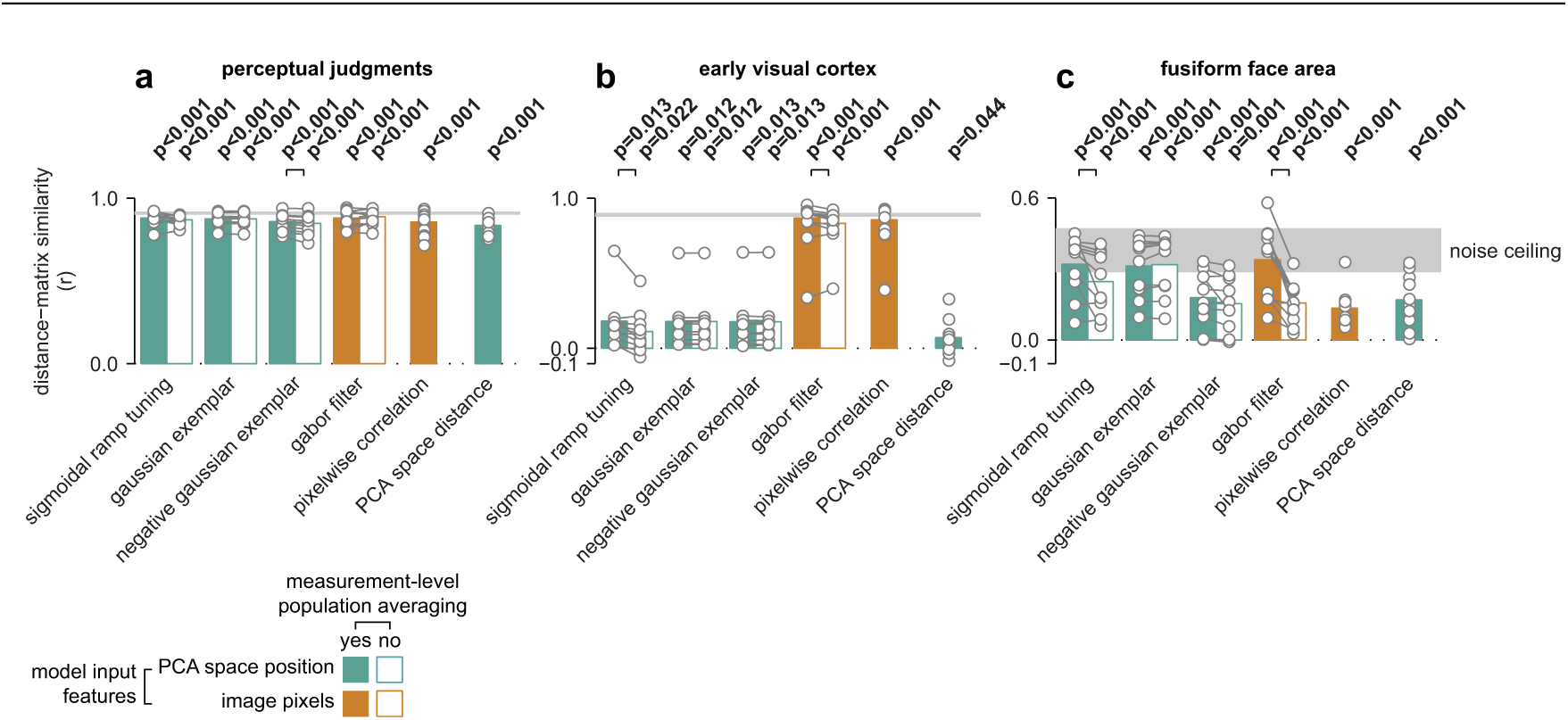
Generalization performance (leave-one-participant-out cross-validation) for fitted computational PCA-space (green bars) and image-based (orange bars) models, as well as mean performance for two fixed predictors (final bars in each panel). The bars provide group-averaged Pearson correlation coefficients, while individual participants are overlaid in gray markers. Filled bars indicate full model fits, while outlined bars indicate model fits excluding measurement-level population averaging. Statistically significant differences between model variants with and without population averaging are illustrated with black connection lines (p < 0.05, see S4 Table for all pairwise comparisons). An estimate of the maximal performance expected given signal-to-noise levels in the sample is illustrated as a shaded noise ceiling (Materials and Methods). Performance is plotted in separate panels for perceptual judgments (**a**), early visual cortex (**b**), and the fusiform face area (**c**). For other regions of interest, see S5 Fig.

**Fig 6.**
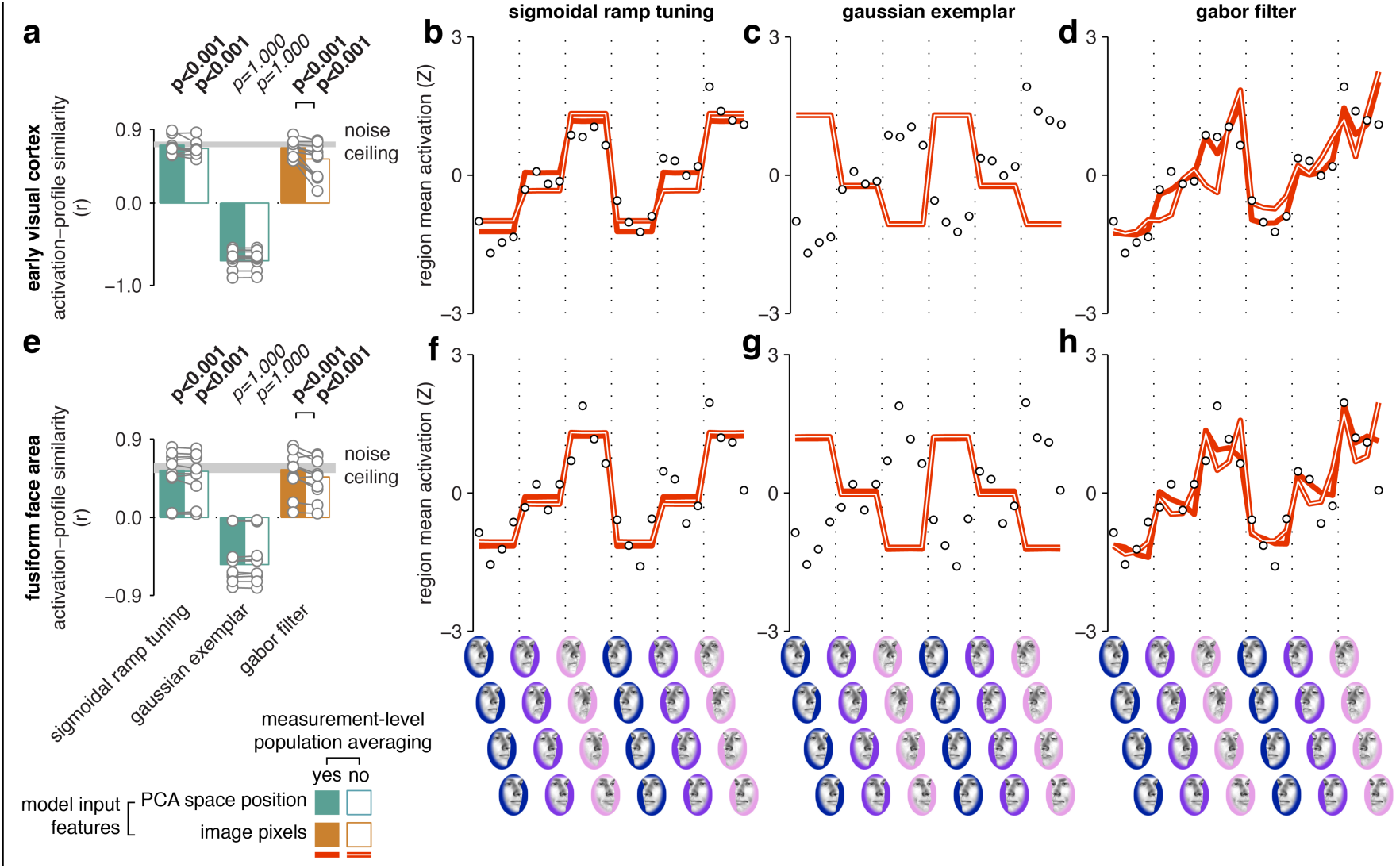
Activation-profile similarity analysis for computational PCA-space and image processing models. Bars (panels **a**, **e**) are plotted similarly as in Fig 5, with group-average effects in bars and single-participant estimates in markers. Markers in the line graphs (panels **b-d, f-h**) indicate group-average fMRI responses evoked by each face exemplar, while lines indicate the computational model predictions (with measurement-level population averaging in filled lines, without in outlined lines). Data and model predictions have been Z-scored to better illustrate the correlation-based fit metric we use here. The top panels illustrate effects in the early visual cortex, while the bottom panels illustrate effects in the fusiform face area. For other regions of interest, see S6 Fig.

### Multiple models can explain cortical and perceptual face spaces

We fitted each of the computational models so as to best predict the representational distance matrices from cortical regions and perceptual judgments. We used a leave-one-participant-out cross-validation approach, in which model performance was evaluated on participants and face identities not used in fitting the parameters (Materials and Methods). Model performance was summarized as the Fisher-Z-transformed Pearson correlation coefficient between the model distances and the data distances. We performed statistical inference on the average Fisher-Z-transformed correlations over all train-test splits of the data using t tests. Our cross-validation scheme tests for generalization across participants and face identities and ensures that models that differ in complexity (number of free parameters) can be compared. In order to investigate the effect of local averaging in fMRI voxels on representational similarity, we fitted two variants of each model: the full model and a variant that excluded measurement-level population averaging.

We found that all evaluated models explained almost all the explainable variance for the perceptual face space (Fig 5a), with only negligible differences in cross-validated generalization performance (for all pairwise model comparisons, see S4 Table). The inclusion of measurement-level population averaging had little effect on performance. This is expected because perceptual judgments, unlike fMRI voxels, are not affected by local averaging across representational units. Thus, the behavioral data was ambiguous with regard to the proposed models, which motivates model selection by comparison to the functional imaging data.

Unlike the perceptual judgments data, the cortical face spaces exhibited substantial differences between the model fits, with a robust advantage for measurement-level population averaging in most cases. In the following, we focus on generalization performance for fits to the early visual cortex and the fusiform face area (for fits to other regions, see S5 Fig).

The early visual cortex was best explained by the Gabor-filter model, which beat the alternative computational models (p < 0.001 for all pairwise model comparisons, S4 Table) and came close to explaining all explainable variance given noise levels in the data. Generalization performance for this model was slightly, but significantly better (p = 0.009, Fig 5b) when population averaging was enabled. As a control, we also tested raw pixel intensities as the representational units. We found no significant difference in performance between the 0-parameter pixel-intensity model and the fitted Gabor-filter model with population averaging (p = 0.380).

The fusiform face area was also well explained by the Gabor-filter model, but in this region we observed similar performance for the sigmoidal-ramp-tuning model. Both models, with population averaging, reached the lower bound of the noise ceiling (Materials and Methods; Fig 5c, S4 Table), suggesting that these models were able to explain the variance in the dataset that was consistent between participants. We also observed comparable generalization performance for the Gaussian exemplar model. Measurement-level population averaging improved generalization performance in the fusiform face area for both the Gabor-filter (p < 0.001) and sigmoidal-ramp-tuning models (p = 0.032), but did not improve either of the exemplar models (p = 0.263 for Gaussian exemplar, p = 0.096 for negative Gaussian exemplar). In summary, the representation in the fusiform face area could be explained by multiple models, and in most cases measurement-level population averaging improved the quality of the fit.

### Regional-mean activation profiles are consistent with sigmoidal-ramp and Gabor-filter, but not Gaussian-exemplar models

Multiple computational models provided qualitatively similar fits to our cortical data at the distance-matrix level. However, we might still be able to adjudicate between them at the level of regional-mean activation profiles. To this end, we obtained activation-profile predictions from each model by averaging over all model units. We then correlated the predicted population-mean activation profile with the regional-mean fMRI activation profile for each participant and performed an activation profile similarity analysis analogously to the distance matrix similarity analysis above. We found that the activation profiles from the computational models were predictive of cortical activation profiles, even though these models were fitted to distance matrices rather than to regional-mean fMRI responses (Fig 6, for other regions see S6 Fig). In particular, the sigmoidal-ramp and Gabor-filter models both predicted increasing population-mean responses with face-space eccentricity, while the Gaussian-exemplar model predicted decreasing responses with eccentricity (S5 Table, S6 Table for pairwise comparisons). This constitutes evidence against the Gaussian-exemplar model, under the assumption that neuronal activity is positively associated with regional-mean fMRI response in visual regions (29–33). The preference for faces closer to the PCA-space origin (sub-caricatures) in the Gaussian-exemplar model arises as a necessary consequence of the Gaussian distribution of preferred faces, which is centered on the average face. We also tested a Gaussian exemplar-tuning model with an inverted Gaussian distribution of preferred faces. In this model, more units prefer faces far from the norm (caricatures) than faces close to the norm (sub-caricatures). However, the inverted-Gaussian exemplar model’s generalization performance was considerably worse than the standard-Gaussian exemplar model’s (Fig 5).

In summary, exemplar models accurately predicted representational distances for cortical face spaces when the preferred-face distribution was Gaussian, but such distributions led to inaccurate predictions of regional-mean fMRI activation profiles. Thus, analysis of regional-mean fMRI responses enabled us to adjudicate between models that made similar predictions at the distance-matrix level, and specifically indicated that the Gaussian exemplar model is unlikely to be the correct model for cortical face-space representation, despite a good fit at the distance-matrix level.

### Measurement-level population averaging is necessary to account for symmetric-view tolerance and over-representation of eccentricity

We found that models that included measurement-level population averaging generally outperformed models that did not. This advantage appeared to originate in how models with population averaging captured two effects in the cortical face spaces: symmetric view-tolerance and over-representation of face-space eccentricity relative to direction.

First, population averaging enabled the image-based Gabor-filter model to exhibit symmetric-view-tolerant responses to the face exemplars. Two mirror-symmetric views will drive a mirror symmetric set of Gabor features. Thus, while the pattern of activity differs, the population-mean activity is similar. Measurement-level pooling over filters centered on distinct visual field locations therefore renders symmetric views more similar in the representation. For instance, the face space in the fusiform face area exhibited little sensitivity to viewpoint, and the Gabor-filter model fit to this region was greatly improved by the inclusion of measurement-level population averaging (Fig 5c, S2 Fig). Indeed, with population averaging, generalization performance for the image-based Gabor-filter model was similar to the sigmoidal-ramp-tuning and exemplar models, for which view tolerance is assumed at the input stage. General view tolerance, beyond the symmetric views we used here, is computationally more challenging. However, for our stimulus set, it was not necessary to posit any intrinsic view-invariant computations in the fusiform face area to explain how its face spaces come to exhibit symmetric view tolerance.

Second, measurement-level population averaging increased the degree to which both the sigmoidal-ramp and the Gabor-filter model over-represented face-space eccentricity relative to direction, which improved the fit for multiple cortical regions. To isolate this smaller effect from the larger symmetric-view-tolerance effect, we collapsed viewpoint in the first-level single-participant fMRI linear model and re-estimated the cortical face spaces and all model fits for the resulting simplified 12-condition design matrix, where each predictor coded appearances of a given face identity regardless of its viewpoint (S7 Fig). Even after collapsing across viewpoints at the first level in this way, measurement-level population averaging still improved the generalization performance of the sigmoidal-ramp and the Gabor-filter model in nearly all cases (p < 0.05, see S7 Table for descriptive statistics and S8 Table for all pairwise comparisons), including the early visual cortex (p < 0.001 for sigmoidal ramp tuning, p = 0.004 for Gabor filter) and the fusiform face area (p = 0.001 for sigmoidal ramp tuning, p = 0.007 for Gabor filter). Thus, the advantage for measurement-level population averaging could not be accounted for by the fact that it helps explain symmetric view tolerance. In sum, the addition of population averaging to the model improved model generalization performance, and this advantage appeared to originate in accounts for two distinct observed phenomena.

## Discussion

This study investigated human face processing by measuring how a face space of individual exemplars was encoded in visual cortical responses measured with fMRI and in perceptual judgments. Relative to a reference PCA model of the 3D shape and texture of faces, cortical face spaces from all targeted regions systematically over-represented eccentricity relative to direction (i.e., the radial relative to the tangential axis). Cortical regions varied in their sensitivity to face viewpoint. We fitted multiple computational models to the data. Considered collectively, the cortical face spaces in the fusiform face area were most consistent with a PCA-space-based sigmoidal-ramp-tuning model and an image-based Gabor-filter model, and less consistent with models based on exemplar coding. As expected, effects in the early visual cortex were consistent primarily with the Gabor-filter model. In all cases, the winning models’ performance depended on the inclusion of a measurement-level population-averaging mechanism, which approximates how individual model units are locally averaged in functional imaging measurements.

### Functional MRI responses in the fusiform face area are best explained by sigmoidal-ramp-tuning and Gabor-filter models

Out of the models we considered, the best accounts for the fusiform face area were a PCA-space-based sigmoidal-ramp-tuning model and an image-based Gabor-filter model. Exemplar-coding models exhibited relatively lower generalization performance, or made inaccurate predictions for regional-mean fMRI activation profiles. Importantly, the advantage for both the sigmoidal-ramp and Gabor-filter model depended on the measurement-level population averaging mechanism. The key contribution of our modeling effort is to narrow the set of plausible representational models for the fusiform face area to two models that can explain both representational distances and the regional-mean activation profile.

It may appear surprising that a PCA-space coding model based on sigmoidal-ramp tuning and an image-based model based on Gabor filters should perform so similarly when fitted to face spaces in the fusiform face area. However, the sigmoidal-ramp-tuning model captures continuous variation in face shape and texture, which covaries with low-level image similarity. For instance, local curvature likely increases with face space eccentricity and is encoded in a ramp-like manner at intermediate stages of visual processing in macaque V4 (27). Conversely, the Gabor-filter model likely possesses sensitivity to face-space direction because the contrast of local orientation content varies with major face features such as eyebrow or lip thickness. Similarly, the Gabor-filter model’s ability to account for regional-mean activation profiles likely arises because local contrast increases with face-space eccentricity, which results in an overall greater activation over the filter banks. There are multiple ways to parameterize face space, not all of which require domain-specific face features. This might also clarify why scene-selective areas such as the parahippocampal place area exhibited somewhat similar representational spaces as face-selective regions in the current study. Such widely-distributed face-exemplar effects are consistent with previous decoding studies (2,4). The Gabor-filter model provides one simple account for how such widely distributed face-exemplar effects can arise. It is likely that the face-space effects we report are driven at least in part by a general mechanism for object individuation in visual cortex rather than the engagement of specialized processing for face recognition.

### Symmetric-view tolerance and over-representation of eccentricity in representational distances can be modeled as an fMRI measurement effect

The models we evaluate here raise the provocative possibility that in some cases, fMRI effects that might conventionally be attributed to high-level featural coding could instead arise from the neuroimaging measurement process. Such an explanation appears possible for two effects in our data. First, even though the Gabor-filter model is a single-layer network with limited representational flexibility, this model nevertheless exhibited near-complete tolerance to mirror-symmetric viewpoint changes, when coupled with measurement-level population averaging. Second, both this model and the sigmoidal-ramp-tuning model showed greater over-representation of eccentricity when measurement-level population averaging was enabled, suggesting that this over-representation in the fMRI data might also plausibly arise through local averaging in voxels.

Previous studies have tended to interpret view-tolerant fMRI effects in terms of cortical processing to support invariant object recognition (2,34,35). The Gabor-filter model suggests a mechanism by which functional imaging measures can exaggerate apparent view-tolerance through spatial pooling over neuronal responses. This result does not contradict previous reports of view-tolerant coding for faces in neuronal population codes measured with single-unit recording (36–39), but rather demonstrates that the type of tolerance to symmetric viewpoint changes that we observed in the current study can be explained without resorting to such intrinsic view-tolerant mechanisms (see also Ramirez et al. (40)). Such findings may go some way toward reconciling apparent discrepancies between single-unit and functional imaging data. For instance, tolerance to symmetrical viewpoints is widespread in human visual cortex when measured with fMRI (34), but appears specific to a subset of regions in the macaque face-patch system when measured with single-unit recordings (36). These results are only contradictory if the measurement process is not considered. In summary, we demonstrate that measurement effects can produce apparent view tolerance in fMRI data. This finding does not suggest that fMRI cannot detect view-tolerant coding (see also 41). For example, population averaging may not account for all cases of non-symmetric view tolerance. However, our results do suggest that modeling of the measurement process is important to correctly infer the presence of such mechanisms from neuroimaging measurements.

### Greater regional-mean activation for distinctive faces can arise from local averaging of neuronal responses

The winning models in this study exemplify how sensitivity to face-space eccentricity at the regional-mean activation level can arise as an artifact of averaging, with no individual neuron encoding distinctiveness or an associated psychological construct. Previous functional imaging studies often interpreted response modulations with face eccentricity as evidence for coding of distinctiveness or related social perception attributes (21–23,42). However, both the PCA-space-based sigmoidal-ramp-tuning model and the image-based Gabor-filter model exhibited increasing population-average responses with eccentricity, even though neither model encodes eccentricity at the level of its units. Although one could, of course, construct a competing model that explicitly codes eccentricity, the models used here are more consistent with single-unit recording studies, where cells generally are tuned to particular features, with a preference for extreme values, rather than responding to eccentricity regardless of direction (20,27,28). Here we demonstrate that when the local averaging of such biologically plausible neuronal tunings is modeled, eccentricity sensitivity emerges without specialized encoding of this particular variable. Related effects have been reported in attention research, where response-gain and contrast-modulation effects at the single-neuron level may sum to similar additive-offset effects at the fMRI-response level (43). In summary, direct interpretation of regional-mean fMRI activations in terms of neuronal tuning can be misleading when the underlying neuronal populations are heterogeneous.

### Modeling of measurement-level population averaging is important for computational studies of cortical representation

A simple model of measurement-level population averaging was sufficient here to substantially improve the generalization performance of multiple computational models for multiple cortical regions. The precise way that fMRI voxels sample neuronal activity patterns remains a topic of debate (30,31,33,44). However, under the simple assumption that voxels sample random subsets of neurons by non-negatively weighted averaging, the effect on the measured fMRI distance matrix will be a uniform stretching along the all-one vector (representing the neuronal population average). To appreciate this point, consider the case of measurement channels that are unlike fMRI voxels in that they sample with random positive and negative weights. Under such conditions, we expect neuronal distances to be approximately preserved in the measurement channels according to the Johnson-Lindenstrauss lemma (for further discussion of this point, see) (45). Intuitively, the measurement channels re-describe the space with randomly oriented axes (without any directional bias). However, fMRI voxels are better understood as taking local non-negative weighted averages of neuronal activity, since the association between neural responses and fMRI response is generally thought to be positive. In such representational spaces the axes have orientations that are biased to fall along the all-1 vector. In practice, this measurement model assumes that the fMRI distance matrix for a given region of interest will over-represent distances to the extent that those distances modulate the neuronal population-average response.

Here we approximated measurement effects for models with nonlinear parameters by mixing the population average into the predicted representational feature space. Despite the simplicity of this method, our noise-ceiling estimates indicate that the winning models captured nearly all the explainable variance in the current dataset. For model representations without nonlinear parameters, this measurement model can be implemented more easily by linearly combining the model’s original distance matrix and the distance matrix obtained for the population average dimension of the space (using squared Euclidean distances estimates, see also 46–48). Thus, the measurement-level population averaging mechanism we propose here is widely and easily applicable to any case where a computational model is compared to neuroimaging data at the distance-matrix level.

The distance-matrix effects of local pooling of neuronal responses in fMRI voxels is correctly accounted for by our measurement model under the assumption that neurons are randomly intermixed in cortex (i.e. voxels sample random subsets of neurons). This simplifying assumption is problematic for early visual areas, where there is a well-established retinotopic organization with a strong contralateral response preference. For the retinotopic Gabor filter model, we therefore added a hemifield-specific pooling stage, which helped account for strong view-sensitivity in occipital regions of interest. Accounting for measurement effects in the presence of topographic organization is likely to prove more challenging for naturalistic stimulus sets. One solution is to account for local averaging in fMRI voxels by local averaging of the model’s internal representational map (45). This local-pooling approach can be thought of as providing a further constraint on the comparison between model and data, because smoothing the model representation is only expected to improve the fit if the model response topography resembles the cortical topography. This may provide a means of adjudicating between topographically organized models, even when the models predict similar distance matrices in the absence of measurement-effect modeling. In summary, the global population average is a special dimension of the representational space, which is overrepresented in voxels that pool random subsets of neurons. This effect accounts for much variance in the representational distances in the current study, and is easy to model. Modeling the overrepresentation of the global average, as we did here, is suitable for models that do not predict a spatial organization (e.g. PCA-space coding models). For models that do predict a spatial organization, it may be more appropriate to simulate fMRI voxels by local averaging of the model’s representational map (45).

### Model comparison is essential for computational neuroimaging

This study demonstrates the importance of considering multiple alternative models to guide progress in computational neuroimaging. In particular, the finding that practically every model we evaluated exhibited significantly greater-than-zero generalization performance strongly suggests how studies that only evaluate a limited set of candidate models can arrive at misleading conclusions (see e.g. 49).

Representational similarity analysis has two key advantages for model comparison relative to alternative approaches. First, competing model predictions can be easily visualized at the level of the best-fitting distance matrix for a given cortical region (e.g., S2 Fig). By contrast, models that are fitted to individual voxels are harder to visualize because the number of voxels per region typically exceeds what can be practically plotted. Furthermore, individual time-points in a rapid event-related fMRI experiment cannot easily be labeled according to experimental stimuli or conditions, which complicates interpretation of fitted time-courses. Second, RSA makes it possible to compare model fits and estimated parameters across data modalities, for instance, between fMRI responses and psychophysical similarity judgments. Such comparisons are challenging when the data is modeled at a lower level, because modality-specific parameters must be added to each model (e.g., parameters controlling the hemodynamic response function for fMRI, decision-threshold parameters for behavioral judgments). The presence of these non-shared parameters makes it difficult to attribute any apparent modality differences to the data rather than to the model specification. In summary, RSA is a particularly attractive analysis approach for studies that emphasize model comparison.

Although the central goal of model comparison is to select the best account of the data, the finding that some models are not dissociable under the current experimental context also has important implications for the design of future studies. Here we demonstrated that Gaussian-distributed exemplar-coding models are less likely to account for human face coding, while accounts based on sigmoidal ramp tuning and Gabor filter outputs perform very similarly. This suggests the need to design stimulus sets that generate distinct predictions from these winning models. For example, presenting face stimuli on naturalistic textured backgrounds may be sufficient to adjudicate between the two models, because the Gabor-filter model lacks a mechanism for figure-ground separation. In conclusion, our study exemplifies the need to test and compare multiple models and suggests routes by which the sigmoidal-ramp-tuning model of face-space coding could be further evaluated.

## Materials and Methods

### Ethics Statement

All procedures were performed under a protocol approved by the Cambridge Psychology Research Ethics Committee (CPREC). Human participants provided written informed consent at the beginning of each data recording day.

### Participants

10 healthy human participants participated in a similarity judgment task and fMRI scans. The psychophysical task comprised 4 separate days of data collection which were completed prior to 4 separate days of fMRI scans. Participants were recruited from the local area (Cambridge, UK) and were naïve with regard to the purposes of the study. Five additional participants participated in data collection up to the first MRI data recording day, but were not invited to complete the study due to difficulties with vigilance, fixation stability, claustrophobia and/or head movements inside the scanner. The analyses reported here include all complete datasets that were collected for the study.

### Data and software availability

The distance matrices we estimated for cortical and perceptual face spaces are available, along with software to re-generate all computational model fits (separate copies deposited on https://osf.io/5g9rv; https://doi.org/10.5281/zenodo.242666). Raw fMRI data will also be available on openfmri.org in the near future.

### Sampling the reference PCA face space

We generated faces using a norm-based model of 3D face shape and texture, which has been described in detail previously (15,25). Briefly, the model comprises two PCA solutions (each trained on 200 faces), one based on 3D shape estimated from laser scans and another based on texture estimated from digital photographs. The components of each PCA solution are considered dimensions in a space that describes natural variation in facial appearance. All stimulus generation was performed using the PCA solution offered by previous investigators, and no further fitting was performed for this study. We yoked the shape and texture solutions in all subsequent analyses since we did not have distinct hypotheses for these.

We developed a method for sampling faces from the reference PCA space in a manner that would maximize dissimilarity variance. This is related to the concept of design efficiency in univariate general linear modeling (50), and involves maximizing the variance of hypothesized distances over the stimulus set. Because randomly sampled distances in high dimensional spaces tend to fall in a narrow range of distances relative to the norm (51), we reduced each participant’s effective PCA space to 2D by specifying a plane which was centered on the norm of the space and extended at a random orientation. The face exemplars constituted a polar grid on this plane, with 4 directions at 60 degrees separation and 3 eccentricity levels (scaled at 30%, 100% and 170% of the mean eccentricity in the training face set). The resulting half-circle grid on a plane through the high-dimensional space is adequate for addressing our hypotheses concerning the relative role of direction and eccentricity coding under the assumption that the high-dimensional space is isotropic. The use of a half-circle also serves to address a potential concern that apparent eccentricity sensitivity might arise as a consequence of adaptation to the experimental stimuli (52). Such adaptation effects are only collinear with eccentricity (ie, prototype) coding if the prototype is located at the average position over the experienced stimulus images. For our stimulus space, this average position would fall approximately between sub- and typical faces and between the second and third direction in the PCA space. Contrary to an adaptation account, there was no suggestion in our data that cortical distances were exaggerated as a function of distance along this axis. The orientation of the PCA-space slice was randomized between participants and model fits were based on cross-validation over participants. Under these conditions, any non-isotropicity is only expected to impair generalization performance. In preliminary tests we observed that this method yielded substantially greater dissimilarity variance estimates than methods based on Gaussian or uniform sampling of the space.

### Face animation preparation

We used Matlab software to generate a 3D face mesh for each exemplar. This mesh was rendered at each of the orientations of interest in the study in a manner that centered the axis of rotation on the bridge of the nose for each face. This procedure ensured that the eye region remained centered on the fixation point throughout each animation in order to discourage eye movements. Renders were performed at sufficient increments to enable 24 frames per second temporal resolution in the resulting animations. Frames were converted to gray-scale and cropped with a feathered oval aperture to standardize the outline of each face and to remove high-contrast mesh edges from the stimulus set. Finally, we performed a frame-by-frame histogram equalization procedure where the average histogram for each frame was imposed on each individual face. Thus, the histogram was allowed to vary across time but not across faces. Note that histogram matching implies that the animations also have identical mean gray-scale intensity and root-mean-square contrast.

A potential concern with these matching procedures is that they could affect the validity of the comparison to the reference PCA space. However, we found that the opposite appeared to be true: distances in the reference PCA space were more predictive of pixelwise correlation distances in the matched images than in the original images. Thus, the matching procedure did not remove features that were encoded in the PCA space and may in fact have acted to emphasize such features.

### Perceptual similarity judgment experiment

We used a pair-of-pairs task to characterize perceptual similarity (S1 Movie). Participants were presented with two vertically offset pairs of faces on a standard LCD monitor under free viewing conditions, and judged which pair was relatively more dissimilar with a button press on a USB keyboard (two-alternative force choice). Each face rotated continuously between a leftward and a rightward orientation (45 degrees left to 45 degrees right of a frontal view over 3 seconds). Ratings across all possible pairings of face pairs (2145 trials: all pairings of the 66 possible pairs of the 12 faces) were combined into a distance matrix for each participant, where each entry reflects the percentage of trials on which that face pair was rated as relatively more dissimilar. The behavioral data was collected over 16 runs (135 trials for the first 15 runs, 120 in the final run). Each participant completed 4 runs in each of 4 data recording days.

### Functional MRI experiment

We measured brain response patterns evoked by faces in a rapid event-related fMRI experiment (S2 Movie). Participants fixated on a central point of the screen where a pseudo-random sequence of face animations appeared (7 degrees visual angle in height, 2s on, 1s fixation interval). We verified fixation accuracy online and offline using an infrared eye tracking system (Sensomotoric Instruments, 50Hz monocular acquisition). The faces rotated outward in leftward and rightward directions on separate trials (18 to 45 degrees rotation left or right of a frontal view), and participants responded with a button press to occasional face repetitions regardless of rotation (one-back task). This served to encourage attention to facial identity rather than to incidental low-level physical features. Consistent with a task strategy based on identity recognition rather than image matching, participants were sensitive to exemplar repetitions within viewpoint (mean d’+−1 standard deviation 2.68+-0.62) and to exemplar repetitions where the viewpoint changed (2.39+-0.52).

The experiment was divided into 16 runs where each run comprised 156 trials bookended by 10s fixation intervals. Each scanner run comprised two experimental runs, which were modeled independently in all subsequent analyses. The data was collected on 4 separate MRI data recording days (2 scanner runs per recording day). The trial order in each run was first-order counterbalanced over the 12 faces using a De Bruijn sequence (53) with 1 additional repetition (diagonal entries in transfer matrix) added to each face in order to make the one-back repetition task more engaging and to increase design efficiency (50). The rotation direction in which each face appeared was randomized separately, since a full 24-stimulus De Bruijn sequence would have been over-long (576 trials). Although the resulting 24-stimulus sequences were not fully counter-balanced, we used an iterative procedure to minimize any inhomogeneity by rejecting rotation direction randomizations that generated off-diagonal values other than 0 and 1 in the 24-condition transfer matrix (that is, each possible stimulus-to-stimulus transfer in the sequence could appear once or not at all). These homogeneous trial sequences served to enhance leave-one-run-out cross-validation performance by minimizing over-fitting to idiosyncratic trial sequence biases in particular runs. We modeled the data from each run with one predictor per face exemplar and viewpoint.

### Magnetic resonance imaging acquisition

Functional and structural images were collected at the MRC Cognition and Brain Sciences Unit (Cambridge, UK) using a 3T Siemens Tim Trio system and a 32-channel head coil. There were 4 separate MRI data recording days for each participant. Each recording day comprised 2 runs of the main experiment followed by 2 runs of the functional localizer experiment. All functional runs used a 3D echoplanar imaging sequence (2mm isotropic voxels, 30 axial slices, 192 × 192mm field of view, 128 × 128 matrix, TR = 53ms, TE = 30ms, 15° flip angle, effective acquisition time 1.06s per volume) with GRAPPA acceleration (acceleration factor 2 × 2, 40 × 40 PE lines). Each participant’s functional dataset (7376 volumes over 8 scanner runs for the main experiment) was converted to NIFTI format and realigned to the mean of the first recording day’s first experimental run using standard functionality in SPM8 (fil.ion.ucl.ac.uk/spm/software/spm8/). A structural T1-weighted volume was collected in the first recording day using a multi-echo MPRAGE sequence (1mm isotropic voxels)(54). The structural image was de-noised using previously described methods (55), and the realigned functional dataset’s header was co-registered with the header of the structural volume using SPM8 functionality. The structural image was then skull-stripped using the FSL brain extraction tool (fmrib.ox.ac.uk/fsl), and a re-sliced version of the resulting brain mask was applied to the fMRI dataset to remove artifacts from non-brain tissue. We constructed design matrices for each experimental run by convolving the onsets of experimental events with the SPM8 canonical hemodynamic response function. Slow temporal drifts in MR signal were removed by projecting out the contribution of a set of nuisance trend regressors (polynomials of degrees 0-4) from the design matrix and the fMRI data in each run.

### Cross-validated discriminant analysis

We estimated the neural discriminability of each face pair for each region of interest using a cross-validated version of the Mahalanobis distance (56). This analysis improves on the related Fisher’s linear discriminant classifier by providing a continuous metric of discriminability without ceiling effects. Similarly to the linear discriminant, classifier weights were estimated as the contrast between each condition pair multiplied by the inverse of the covariance matrix of the residual time courses, which was estimated using a sparse prior (57). This discriminant was estimated separately for the concatenated design matrix and fMRI data in each possible leave-one-out split of the experimental runs, and the resulting weights were transformed to unit length and projected onto the contrast estimates from each training split’s corresponding test run (16 estimates per contrast). The 16 run-specific distance estimates were averaged to obtain the final neural discriminability estimate for that participant and region. When the same data is used to estimate the discriminant and evaluate its performance, this algorithm returns the Mahalanobis distance, provided that a full rather than sparse covariance estimator is used (56). However, unlike a true distance measure, the cross-validated version that we use here is centered on 0 under the null hypothesis. This motivates summary-statistic random-effects inference for above-chance performance using conventional T tests.

We developed a variant of this discriminant analysis where effects that might be broadly described as region-mean-related are removed (S4 Fig). This control analysis involved two modifications to how contrasts were calculated at the level of forming the discriminant and at the level of evaluating the discriminant on independent data. First, each parameter estimate was set to a mean of zero in order to remove any additive offsets in response levels between the conditions. Second, for each pair of mean-subtracted parameter estimates, the linear contribution of the mean estimate over the pair was removed from each estimate before calculating the contrast. This corrects for the case where a single response pattern is multiplicatively scaled by the conditions. The resulting control analysis is insensitive to effects driven by additive or multiplicative scaling offsets between the conditions.

### Multiple regression RSA

We used a multiple regression model to estimate the relative contribution of eccentricity and direction to cortical and perceptual face-space representations. Multiple regression fits to distance estimates can be performed after a square transform, since squared distances sum according to the Pythagorean theorem. We partitioned the squared distances in the reference PCA space into variance associated with eccentricity changes by creating a distance matrix where each entry reflected the minimum distance for its eccentricity group in the squared reference PCA matrix (that is, cases along the group’s diagonal where there was no direction change). The direction matrix was then constructed as the difference between the squared reference PCA matrix and the eccentricity matrix (Fig 2a). These predictors were vectorized and entered into a multiple regression model together with a constant term. This partitioning of the dissimilarity variance in the reference PCA matrix is complete in the sense that a multiple regression RSA model where the reference PCA matrix is used as the dependent variable yields parameter estimates of [1,1,0] for eccentricity, direction and constant, respectively, with no residual error. These three predictors were then split according to viewpoint, with separate sets of predictors for distances within and across viewpoint. The absolute values of the cortical and perceptual distance matrices were squared and then transformed back to their original sign before being regressed on the predictor matrix using ordinary least squares. Finally, the absolute values of the resulting parameter estimates were square-root transformed and returned to their original signs.

### Functional regions of interest

We used a conventional block-based functional localizer experiment to identify category-selective and visually-responsive regions of interest in human visual cortex. Participants fixated a central cross on the screen while blocks of full-color images were presented (36 images per block presented with 222ms on, 222ms off, 16 s fixation). Participants were instructed to respond to exact image repetitions within the block. Each run comprised 3 blocks each of faces, scenes, objects and phase-scrambled versions of the scene images. Each participant’s data (8 runs of 380 volumes, 2 runs collected for each of 4 MRI data recording days) was smoothed with a Gaussian kernel (6mm full width at half maximum) and responses to each condition were estimated using a standard SPM8 first-level model. Regions of interest were identified using a region-growing approach, where a peak coordinate for each region was identified in individual participants, and a region of interest was grown as a contiguous set of the most selective 100 voxels extending from this coordinate. We defined the face-selective occipital and fusiform face areas with the minimum-statistic conjunction contrast of faces over objects and faces over baseline, and the scene-selective parahippocampal place area and transverse occipital sulcus as the minimum-statistic conjunction contrast of scenes over objects and scenes over baseline, and the early visual cortex as the contrast of scrambled stimuli over the fixation baseline. We also attempted to localize a face-selective region in the posterior superior temporal sulcus, a face-selective region in anterior inferotemporal cortex and a scene-selective region in retrosplenial cortex, but do not report results for these regions here since they could only be identified in a minority of the participants. All regions of interest were combined into bilateral versions before further analysis since we did not have distinct predictions concerning functional lateralization. Example regions of interest can be viewed in S8 Fig. Typical MNI coordinates for each region are provided in S9 Table.

### Sigmoidal ramp tuning model

The sigmoidal ramp model comprises 1000 model units, each of which exhibits a monotonically increasing response in a random direction extending from the origin of the face space (Fig 3). The response *y* at position *x* along the preferred direction is described by the sigmoid

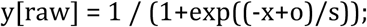

where the free parameters are *o*, which specifies the horizontal offset of the response function (zero places the midpoint of the response function at the norm of the space, values greater than zero corresponds to responses shifted away from the norm), and *s*, which defines response function saturation (4 corresponds to a near-linear response in the domain of the face exemplars used here, while values near zero correspond to a step-like increase in response). The raw output of each model unit is then translated toward the population-mean response

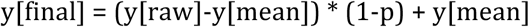

where *p* is a free parameter that defines the strength of measurement-level population averaging (0 corresponds to no averaging, 1 corresponds to each model unit returning the population-mean response).

### Exemplar model

The exemplar model comprises 1000 model units, each of which prefers a Cartesian coordinate in the face space with response fall-off captured by an isotropic Gaussian. The free parameters are *w*, which controls the full width at half-maximum tuning width of the Gaussian response function, and *d*, which controls the width of the Gaussian distribution of tuning centers (0.1 places Z = 2.32 at 10% of the eccentricity of the caricatures while 3 places this tail at 300% of the eccentricity of the caricatures).

We also constructed an inverted-Gaussian variant of this model where the distribution of distances was inverted at Z = 2.32 and negative distances truncated to zero (1% of exemplars). This model was fitted with similar parameters as the original Gaussian exemplar model.

### Gabor filter model

The Gabor filter model is an adaptation of a neuroscientifically-inspired model that has previously been used to successfully predict single-voxel responses in the early visual cortex (24, github.com/kendrickkay/knkutils/tree/master/imageprocessing). The model is composed of 5 banks of Gabor filters varying in spatial position, phase (2 values) and orientation (8 directions, Fig 4). We measured each filter’s response to the last frame of each animation, and corrected for phase shifts by collapsing the two signed phase value filter outputs into a single non-negative estimate of contrast energy (specifically, the square root of the sum over the two squared phase values). This is a standard processing stage in the Gabor filter model (24). The resulting rectified response vectors were weighted according to filter bank membership (5 free parameters). We estimated measurement-level population averaging using two pooling stages: a hemifield-specific pool, where filters were pooled according to whether their centers fell left or right of the vertical meridian, followed by a global pool.

### Pixelwise correlation predictor

We used a fixed control predictor to estimate whether coding based on pixelwise features would produce the same face-space warping we observed in our data (Fig 4). The pixelwise correlation predictor was generated by stacking all the pixels in each of the face animations into vectors and estimating the correlation distance between these intensity values.

### Estimating the noise ceiling

We estimated the noise ceiling for Z-transformed Pearson correlation coefficients based on methods described previously (56). This method estimates the explained variance that is expected for the true model given noise levels in the data. Although the true noise level of the data cannot be estimated, it is possible to approximate its upper and lower bounds in order to produce a range within which the true noise ceiling is expected to reside. The lower bound estimate is obtained by a leave-one-participant-out cross-validation procedure where the mean distance estimates of the training split are correlated against the left-out-participant’s distances, while the upper bound is obtained by performing the same procedure without splitting the data. These estimates were visualized as a shaded region in figures after reversing the Z-transform (Fig 4).

### Statistical inference

All statistical inference was performed using T-tests at the group-average level (N=10 in all cases except the occipital face area and transverse occipital sulcus, N = 9). Correlation coefficients were Z-transformed prior to statistical testing. Average Z statistics were reverse-transformed before visualization for illustrative purposes.

Fold-wise generalization performance estimates are partially dependent, which can lead to sample variance underestimates (58,59) and greater than intended false positive rates when conventional parametric statistics are used. However, we simulated the effects of this potential bias and found no consistent inflation in false-positive rates for simulations of the parameters used in the current study (S1 Code, S9 Fig, S10 Fig). Thus, the inferential statistics reported in the current study appear to be robust to this slight dependence and maintain their intended frequentist properties.

## Acknowledgments

We are grateful to Jenna Parker for assistance with data collection and to Marta Correia for assistance with 3D EPI protocols.

## Supporting information captions

**S1 Fig.**
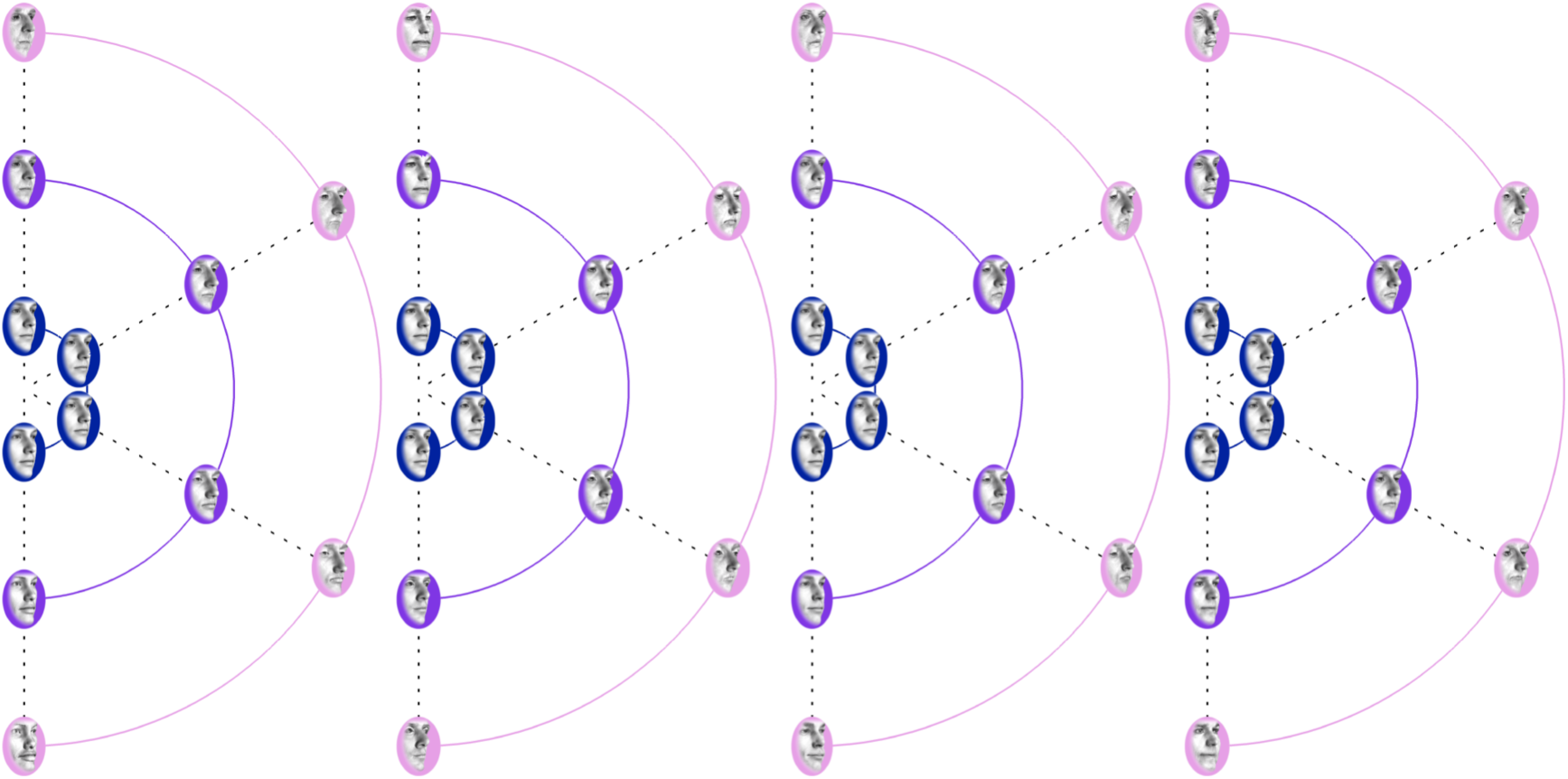
Example stimulus sets for 4 participants. Each stimulus set shares the same underlying distance matrix in the reference PCA space, while the randomization of the orientation of the plane on which the faces are sampled ensures that each set is visually distinct.

**S2 Fig.**
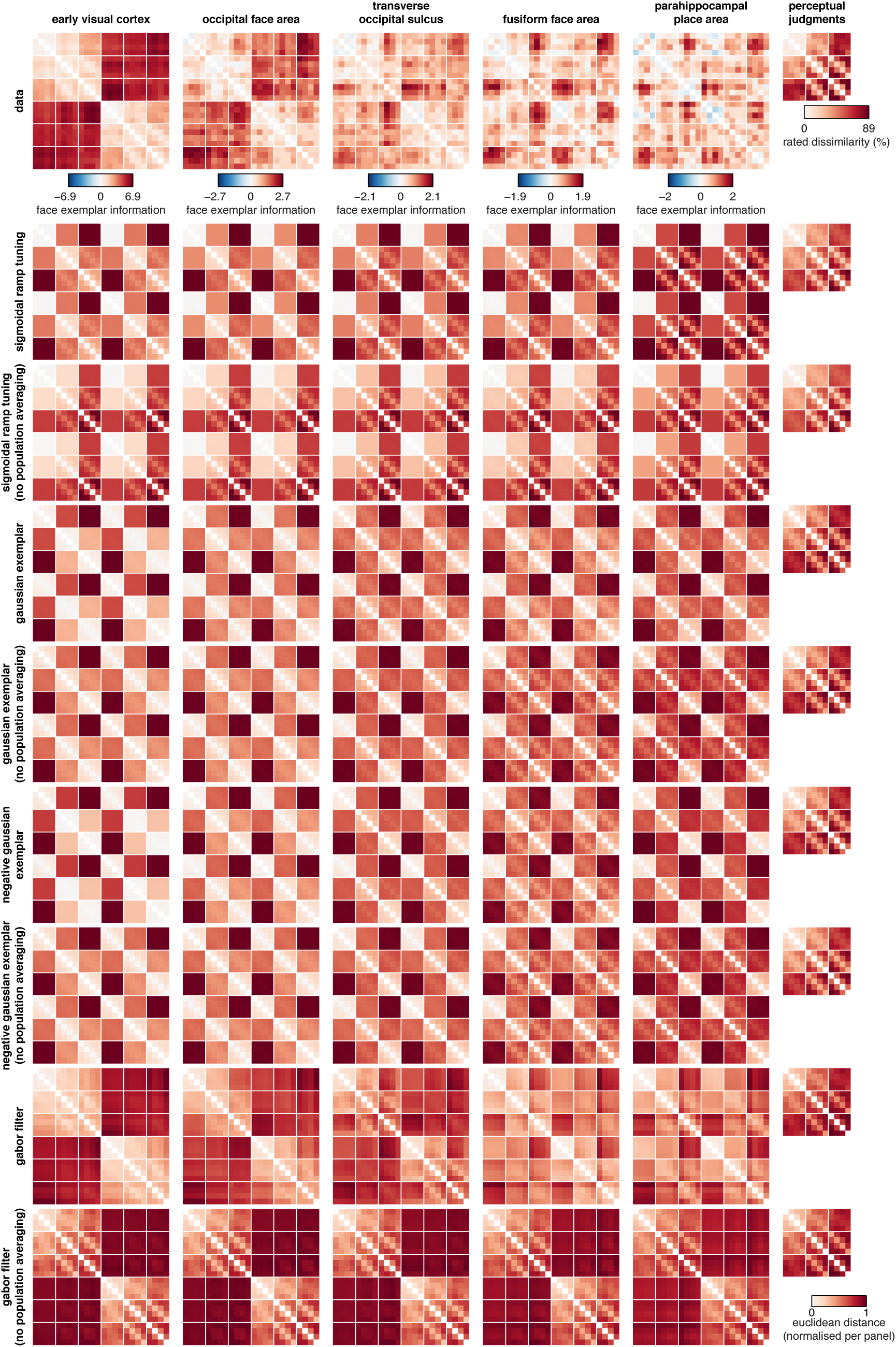
Group-average distance matrices from additional regions of interest and best-fitting model predictions from each model considered in the main manuscript (Fig 5).

**S3 Fig.**
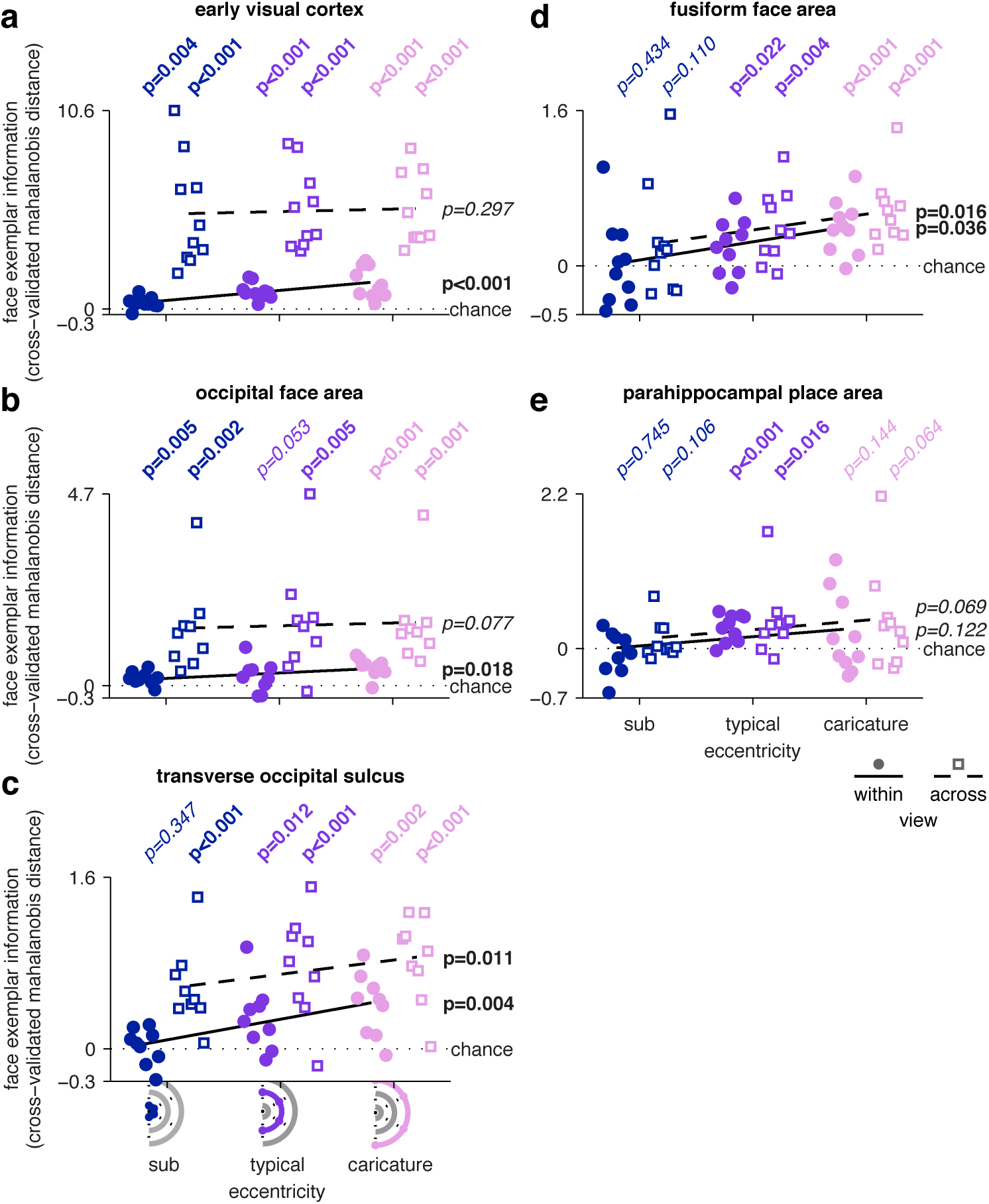
Cortical direction discriminability as a function of eccentricity level. Each point reflects the mean performance for all directions at a given eccentricity level (4 × 4 block diagonals in Fig 1) for a single participant. Small random offsets have been added to each x coordinate for illustrative purposes, and a line shows the least-squares fit. Performance is plotted separately for distances within viewpoint (round markers, solid line, left offset) and across viewpoint (square markers, dashed line, right offset). All plotted p values are obtained through group analysis of single-participant estimates. Within viewpoint, cortical discrimination performance increases with eccentricity level in all regions except the parahippocampal place area (e). Across viewpoint, statistically significant effects are observed in the ventral temporal fusiform face area in the lateral temporal transverse occipital sulcus, but not in occipital areas (early visual cortex, occipital face area).

**S4 Fig.**
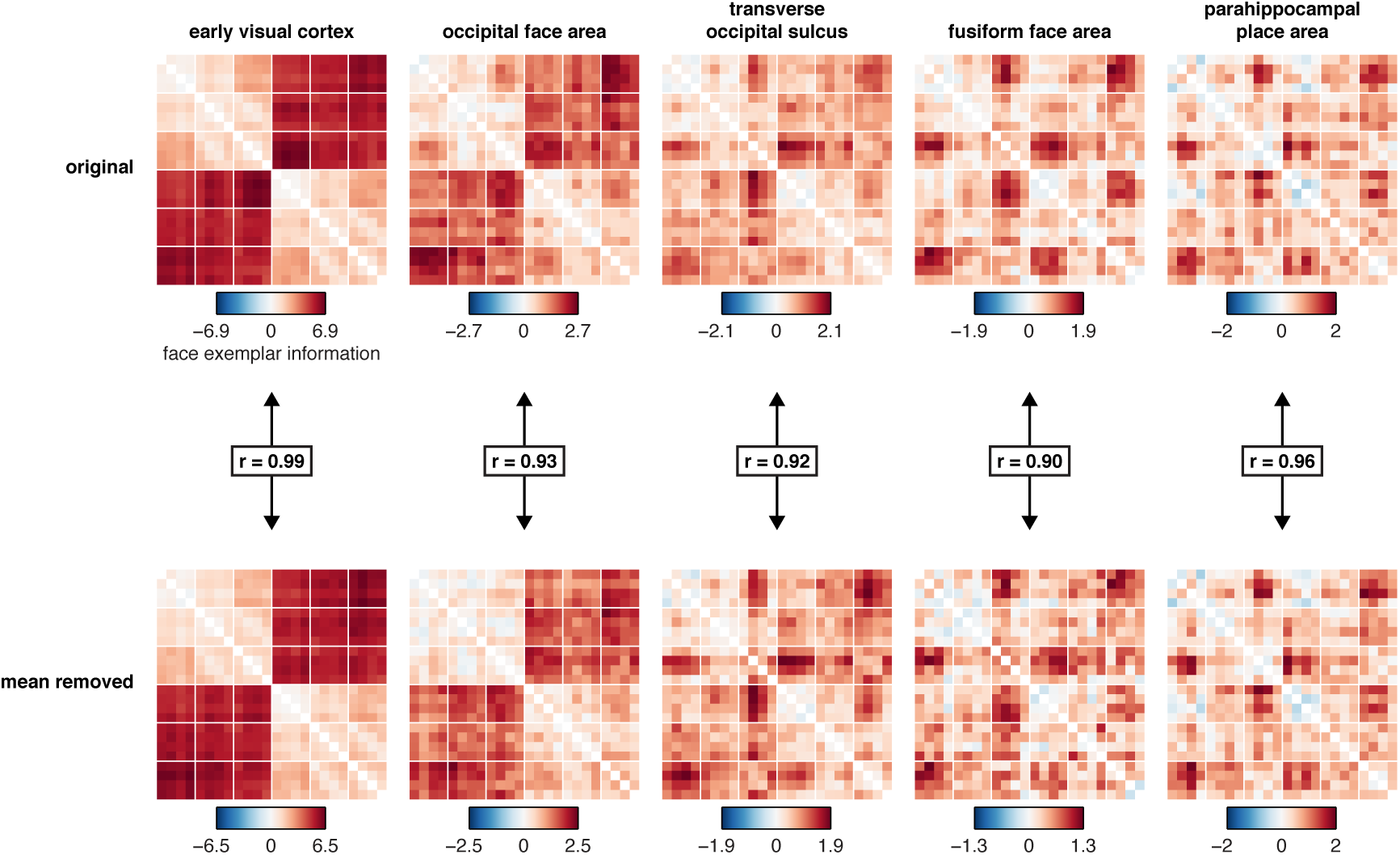
Effect of region-mean removal on cortical face spaces. The top row shows original distance matrices, while the bottom row shows distance matrices after removing additive and multiplicative mean pattern effects (Materials and Methods). The cited Pearson correlation coefficients are calculated at the group-average level.

**S5 Fig.**
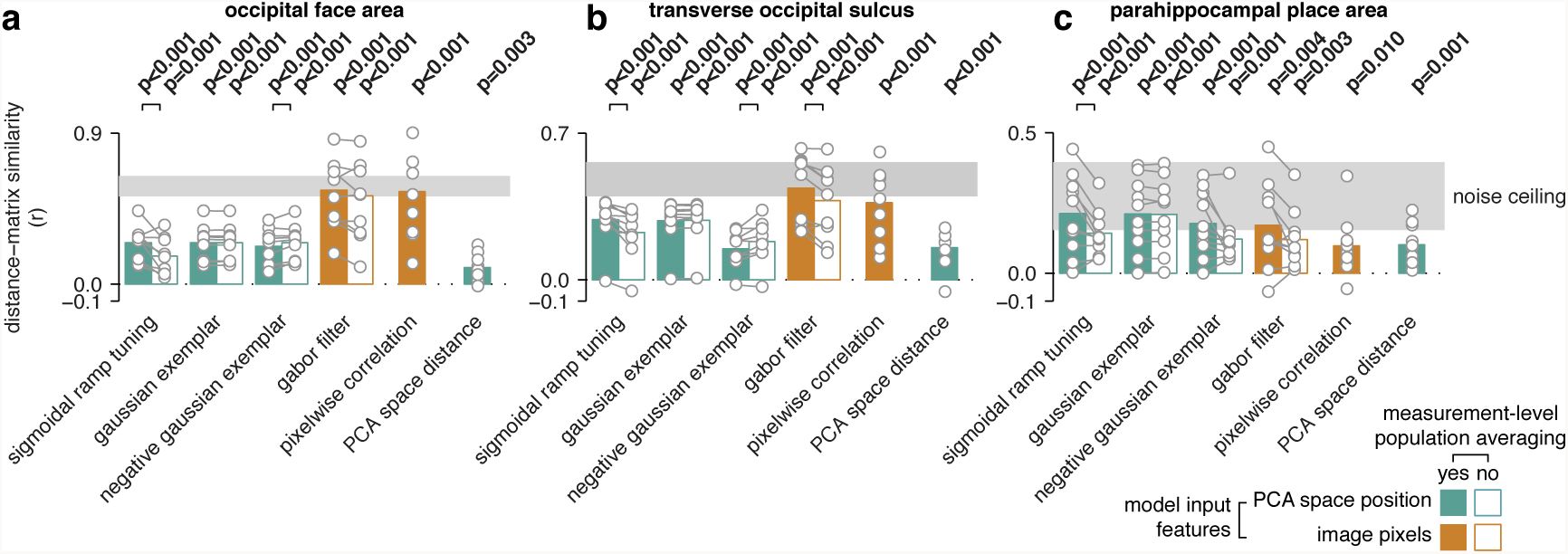
Cross-validated distance-matrix generalization performance for additional cortical regions of interest. Plotted as Fig 5 in main text.

**S6 Fig.**
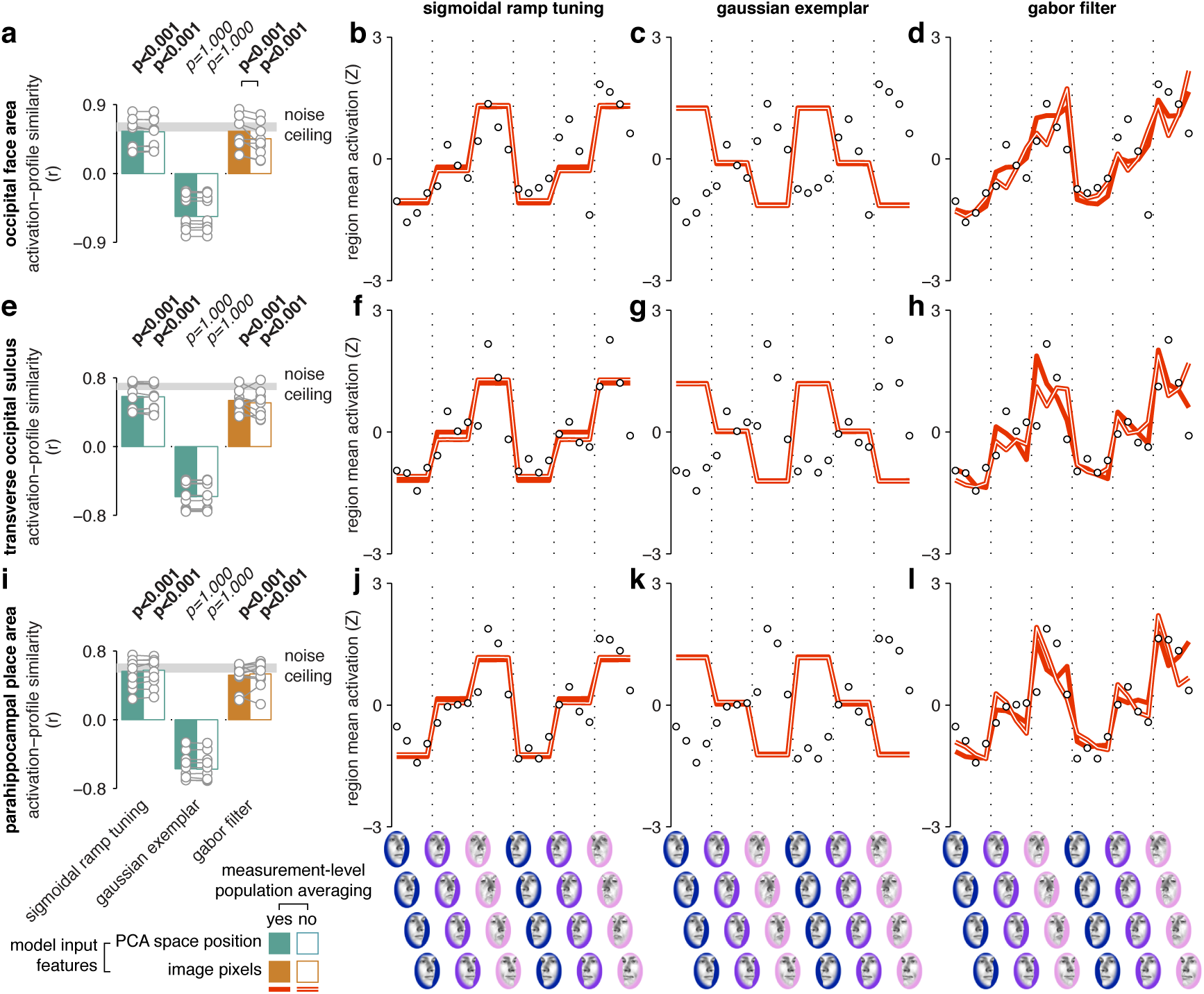
Cross-validated activation-profile generalization performance for additional cortical regions of interest. Plotted as in Fig 6 in main text.

**S7 Fig.**
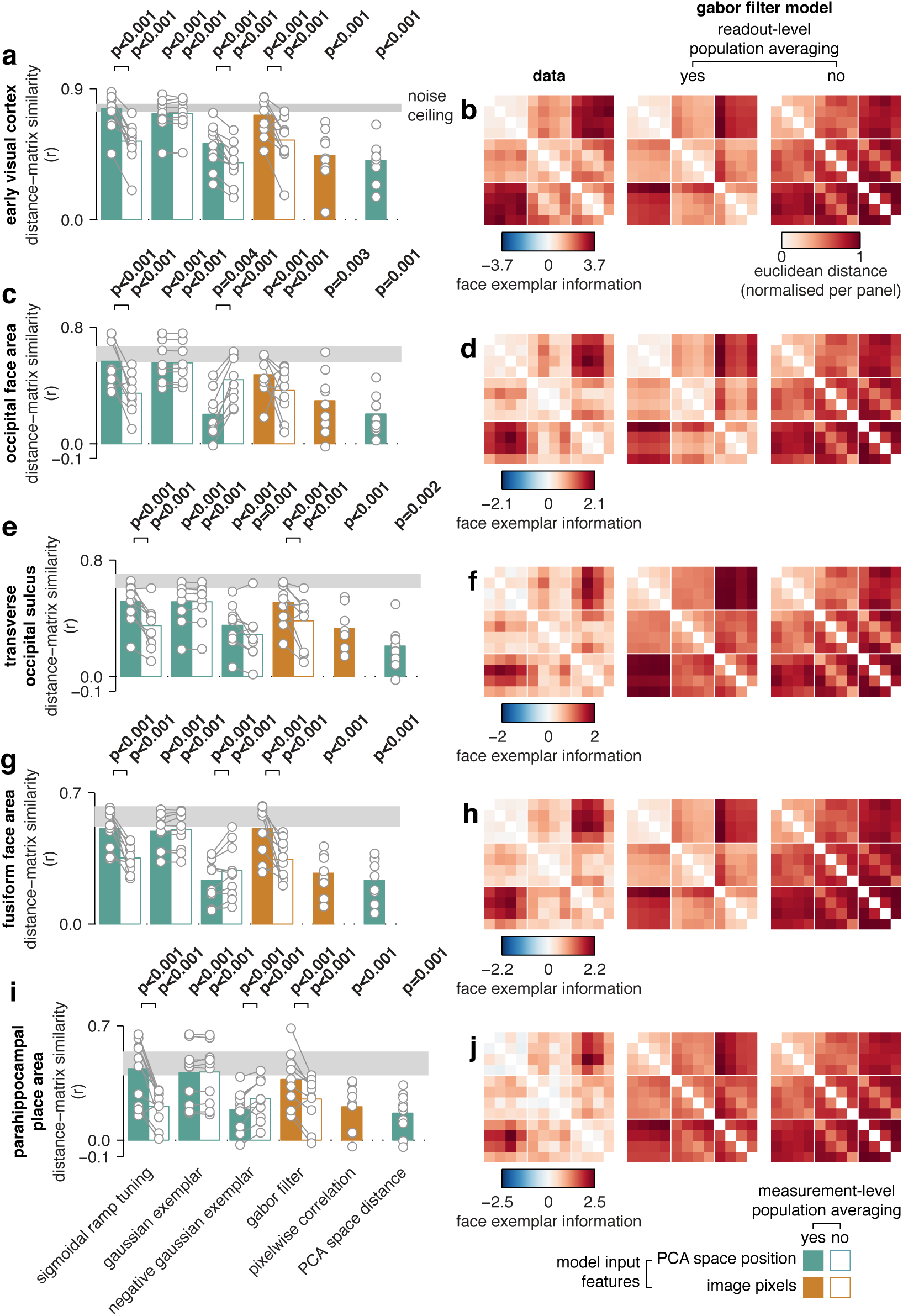
Cross-validated distance-matrix similarity, data distance matrices and best-fitting predicted matrices for an analysis where the two viewpoints have been collapsed into a single set of 12 conditions. Plotted as in Figs 1 and 5 in main text.

**S8 Fig.**
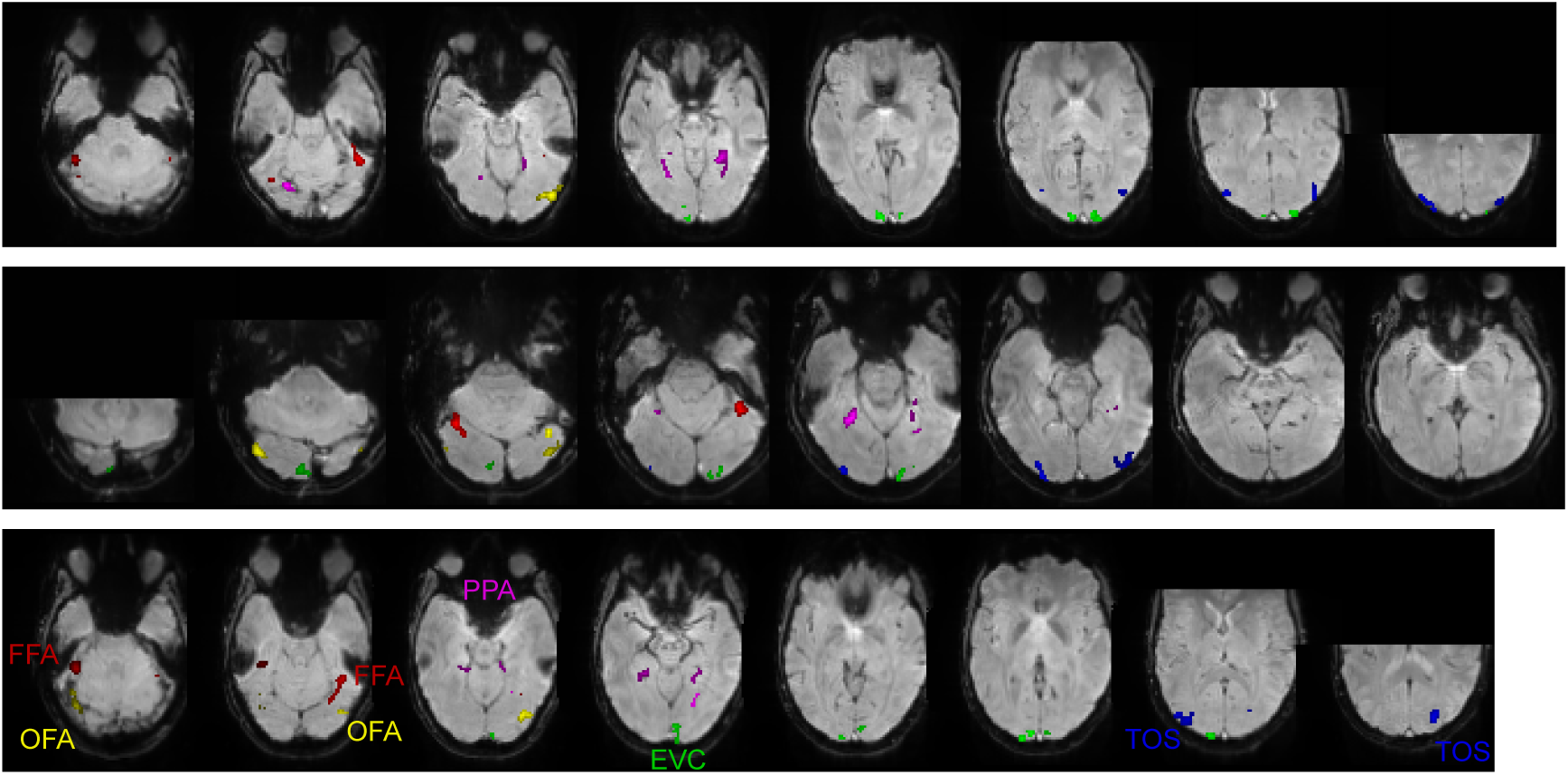
Example cortical regions for 3 participants (rows) overlaid on the mean fMRI volume from the first scanner run of the experiment. Regions are color-coded and labels are provided in the bottom row. Abbreviations and colors: EVC – early visual cortex, green; FFA – fusiform face area, red; OFA – occipital face area, yellow; PPA – parahippocampal place area, purple; TOS – transverse occipital sulcus, blue.

**S9 Fig.**
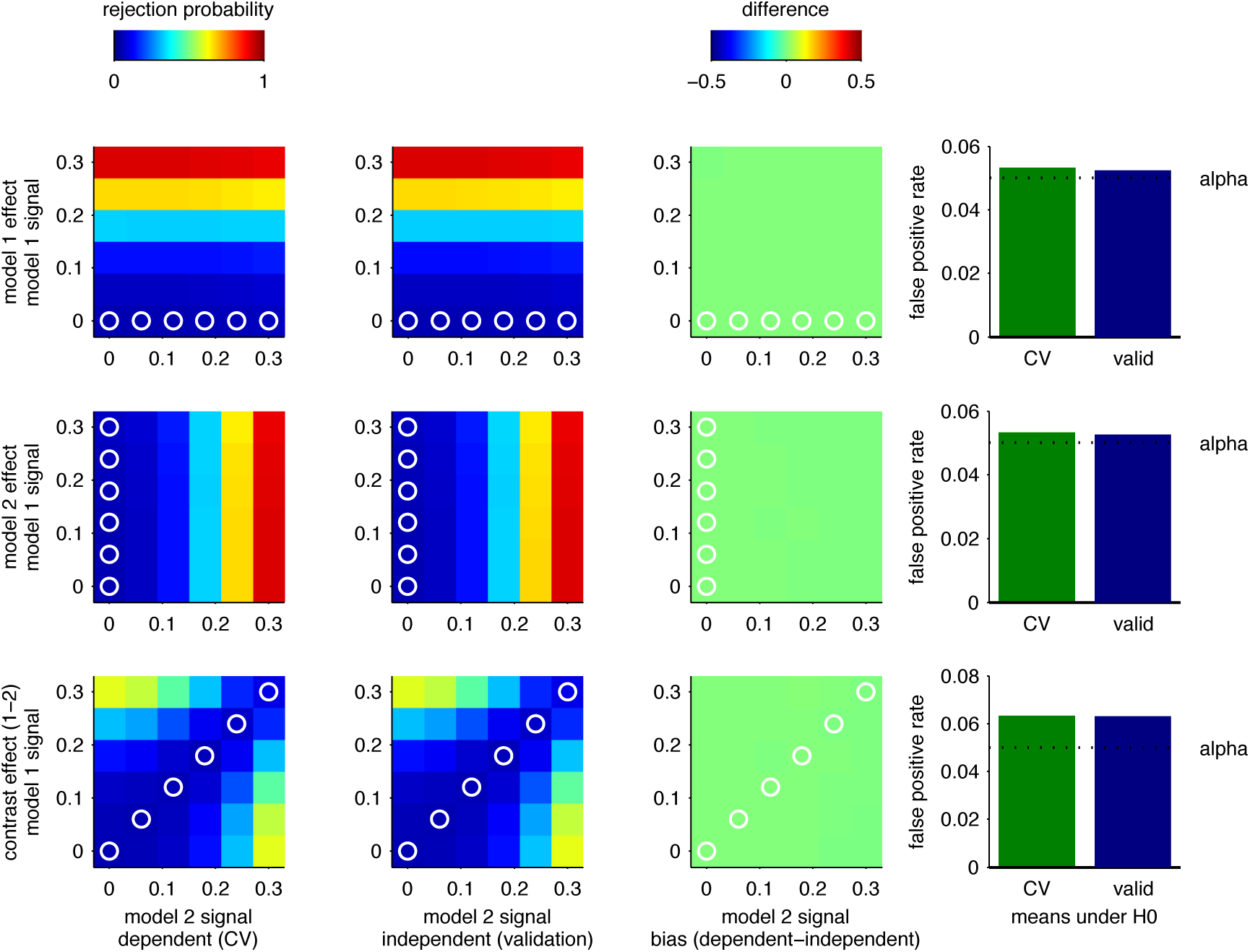
Rejection probability simulation. We simulated out-of-sample generalization performance for two arbitrary models, using methods that closely matched the ones described in this manuscript (for details, see S1 Code). We estimated potential bias by comparing cross-validated generalization performance (panels in leftmost column) with generalization to a withheld validation set (left column, subtraction in right column). The probability of rejecting the null hypothesis (p < 0.05, T test) over 100000 simulations is plotted for tests of either model against zero (one-tailed test, first two rows of panels), and of zero difference between the models’ generalization performance (two-tailed test, bottom row). Each color-mapped image shows the rejection probability as a function of signal level for model 1 (vertical axis) and model 2 (horizontal axis). The null hypothesis case is highlighted with white circles. The bars in the rightmost panel summarize the mean rejection probabilities (ie, false positives) for each of these null cases.

**S10 Fig.**
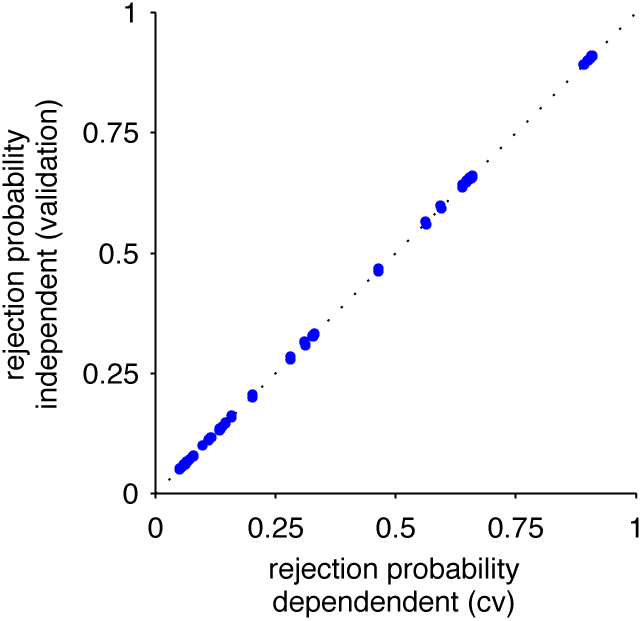
Summary plot of the data in S9 Fig. Each point represents the mean rejection probability for a unique set of simulation parameters (values in color-mapped images in S9 Fig). It can be seen that there is a close to unit relationship between rejection probability in the dependent, cross-validated case (vertical axis) and the independent, validation case (horizontal axis).

*S1 Code.* Code to reproduce S9 Fig and S10 Fig. Requires Matlab R2013a

*S1 Table*. Descriptive and inferential statistics for the distance-matrix correlation between the face-space models and the perceptual and cortical face spaces. We report the group-average correlation coefficient (mean_r), the group-average Z-transformed correlation (mean_zr), standard error for the Z-transformed correlation (sterr_zr), one-tailed p values for the Z-transformed correlation (ppara_zr) and sample sizes (n). Related to Fig 5.

*S2 Table*. Analysis of variance on parameter estimates from multiple regression RSA model, with the factors metric (eccentricity, direction), viewpoint (within, across), and a two-way interaction term. For details, see main text. Related to Fig 2.

*S3 Table*. Descriptive and inferential statistics for the analysis of direction discriminability as a function of eccentricity level. Mean, standard error (sterr), one-tailed p values (ppara) and sample sizes (n) are included on separate rows. See also S3 Fig.

*S4 Table*. Two-tailed parametric p values for all pairwise comparisons between model distance-matrix generalization performances (S1 Table). Related to Fig 5.

*S5 Table*. Descriptive and inferential statistics for activation-profile similarity analysis. See S1 Table for an account of what the row labels represent. Related to Fig 6.

*S6 Table*. Two-tailed parametric p values for all pairwise comparisons between activation-profile model fits (S5 Table). Related to Fig 6.

*S7 Table*. Descriptive and inferential statistics for collapsed-view distance-matrix generalization performances. See S1 Table for an account of what the row labels represent. Related to S7 Fig.

*S8 Table*. Two-tailed parametric p values for all pairwise comparisons between collapsed-view distance-matrix generalization performances. See S7 Table, S7 Fig.

*S9 Table*. Typical coordinates for the cortical regions of interest. We identified the peak voxel in each region of interest using the region-defining contrast in the localizer experiment, and converted these native-space voxel indices to standard mm coordinates in the MNI template brain using transformations obtained from SPM8 structural T1 normalization routines. We report means and standard deviations across participants for each region. See also S8 Fig.

*S1 Movie*. Cropped screen capture of the perceptual judgment task as it appeared to participants during data collection.

*S2 Movie*. Cropped screen capture of the main experiment as it appeared to participants during data collection.

